# Active vision in freely moving marmosets using head-mounted eye tracking

**DOI:** 10.1101/2024.05.11.593707

**Authors:** Vikram Pal Singh, Jingwen Li, Kana Dawson, Jude F. Mitchell, Cory T. Miller

## Abstract

Our understanding of how vision functions as primates actively navigate the real-world is remarkably sparse. As most data have been limited to chaired and typically head-restrained animals, the synergistic interactions of different motor actions/plans inherent to active sensing –e.g. eyes, head, posture, movement, etc.-on visual perception are largely unknown. To address this considerable gap in knowledge, we developed an innovative wireless head-mounted eye tracking system called **CEREBRO** for small mammals, such as marmoset monkeys. Our system performs **C**hair-free **E**ye-**Re**cording using **B**ackpack mounted mic**RO**controllers. Because eye illumination and environment lighting change continuously in natural contexts, we developed a segmentation artificial neural network to perform robust pupil tracking in these conditions. Leveraging this innovative system to investigate active vision, we demonstrate that although freely-moving marmosets exhibit frequent compensatory eye movements equivalent to other primates, including humans, the predictability of the visual behavior (gaze) is higher when animals are freely-moving relative to when they are head-fixed. Moreover, despite increases in eye/head-motion during locomotion, gaze stabilization remains steady because of an increase in VOR gain during locomotion. These results demonstrate the efficient, dynamic visuo-motor mechanisms and related behaviors that enable stable, high-resolution foveal vision in primates as they explore the natural world.

**Significance Statement:** Vision is arguably the most thoroughly understood of all neural systems in the primate brain. Yet there is little known about how vision functions in real-world contexts in which individuals freely move and explore an environment. This dearth in knowledge is largely due to the lack of technology that can accurately track eye-movements in freely-moving individuals with the speed and resolution needed to quantify primate vision. Here we developed an innovative wireless head-mounted eye-tracking system for marmosets that meets these technical needs and enabled us to quantify facts of primate vision in a manner not previously possible, including a set of discoveries that are likely to transform our understanding of this keystone system.

## Introduction

Primate vision has been the subject of intense study for many decades and is arguably the most well understood neural system in the simian brain (1, 2). And yet, our understanding of primate vision is incomplete. Like all sensory systems, primate vision evolved in response to the challenges inherent to actively moving, exploring and engaging with objects, individuals and the environment from different perspectives (3–5). While the processes of visual encoding have been extensively studied with head-restrained subjects observing stimuli presented on a screen, details of how vision functions as primates actively move through and explore the real-world are remarkably limited. Similar to all vertebrates, the head and eyes coordinate their respective movements in primates to enable a stable percept of visual inputs (6–10). Previous studies in chaired but head-free macaques demonstrate how the eye and head coordinate to stabilize gaze but have yet to address similar coordination in freely-moving primates. And in marmosets, though we have some understanding that their oculomotor range is more restricted than macaques when head-fixed (11) and that their head-position can shift more rapidly than humans or macaques (12), we still lack descriptions of the head-eye coordination for gaze shifts in freely-moving scenarios. Because data on this complement of mechanisms has been limited to chaired animals unable to locomote, the synergistic effects of different motor actions – e.g. eyes, head, posture, movement, etc.-on primate visual perception and cognition during active exploration of the world are almost entirely unknown. The principal bottleneck being technical. Several previous studies have sought to bridge this gap and achieved limited precision for projecting the gaze of free-moving non-human-primates (NHPs) into visual scenes (13–15), but due to technical constraints of these methods, no studies have been able to quantify the stability of visual gaze nor the eye and head dynamics during freely moving active exploration. Here we overcome these obstacles and introduce a method that precisely quantifies eye-movements and can accurately project the gaze of a NHP into scenes as individuals freely explore an environment.

Recent work with mice demonstrates that eye-tracking systems can be miniaturized and mounted to the head in order to provide insight into vision during natural visual behavior (16–19), but these systems are not well suited for comparable studies in primates for at least two critical reasons. First, systems in mice rely on a tether which restricts the 3D mobility of primates. Second, they lack the precision and temporal resolution needed to accurately characterize high-resolution vision of primates. To address this challenge, we developed an innovative head-mounted eye tracking system to enable the study of active, natural visual behaviors, and related neural processes, in freely-moving marmosets. Our system -**CEREBRO**-allows for **C**hair-free **E**ye-**Re**cording using **B**ackpack mounted mic**RO**controllers at a speed and resolution needed to accurately quantify the visual behavior and underlying neural mechanisms of natural, active vision in primates. Using CEREBRO we confirmed that freely-moving marmosets exhibit frequent compensatory eye movements that enable them to stabilize gaze when viewing real world scenes consistent with previous studies in body restrained animals. By using this innovative system, however, we discovered that visual gaze stabilization and predictability was enhanced when the monkeys were moving naturally despite increases in eye/head-motion during locomotion. This suggests that previously unreported synergistic mechanisms for gaze stabilization are not only integral to primate active vision in the real-world but can be enhanced for greater compensation as animals move naturally through their environment.

## Results

### *CEREBRO* is a head mounted eye tracking system for freely moving marmosets

The small body size of marmosets (∼300-400g) necessitated certain design considerations when developing CEREBRO. The first of these decisions is related to the weight and wearability of the system itself. Based on previous experience, the estimated 60g weight of the complete system would not be feasible to be entirely situated on animal’s head without affecting its visual behavior. To resolve this issue, we separated the system into two separate, but integrated hardware submodules: the ***Head-piece*** and the ***Backpack***. The *Head-piece* includes the camera assembly and scaffold fitted to the animals’ head *(fig 1a)*; and the *Backpack* includes the backend electronics for camera synchronizing, image acquisition and local data storage on the custom designed printed circuit boards (PCBs) *(fig1b)*. Both the PCB and the head-piece IR LED are powered using a 600 mAh, Lithium-Polymer (Li-Po) battery that is housed inside the backpack. OV4689 camera modules (SincereFirst, Guangzhou, China) were used to record the eye and the world scene. For more details about camera interface see Materials & Methods (Camera and communication interface).

**Fig 1:**
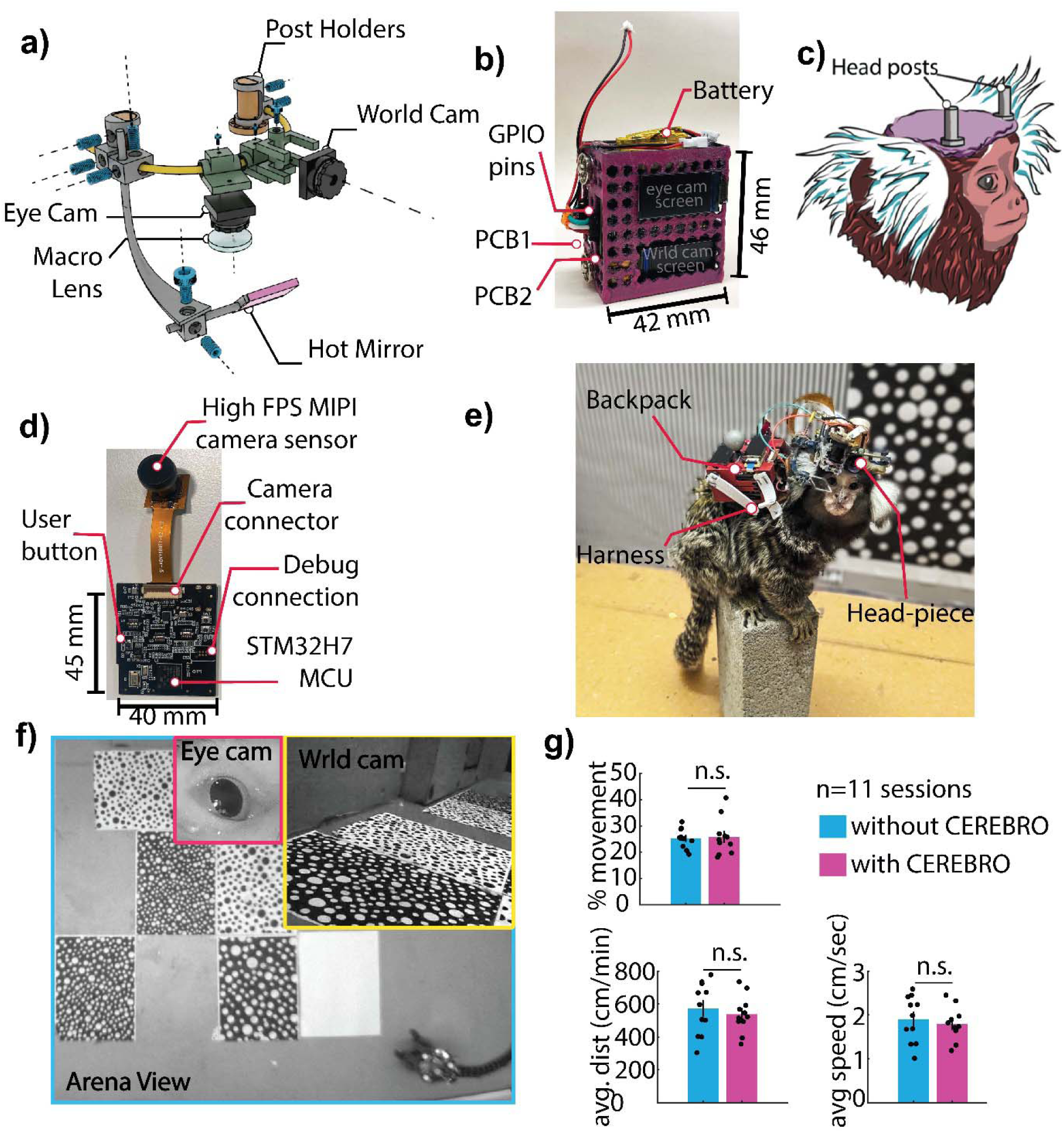
Head mounted eye-tracker assembly. **(a)** (a) 3D render depicting different parts of the head-piece. The Ti-alloy scaffolding is custom designed and fabricated using DLS **(b)** 3D printed backpack that holds the 2 PCBs and the battery. **(c)** The animals are fitted with two 1cm headposts that act as anchors for holding the head-piece. **(d)** Custom PCB board which drives the camera sensors, IMU and saves the data to a locally mounted SD card**. (e)** A marmoset wearing the fully assembled system (CEREBRO). (f) View from the 3 different cameras (eye camera(pink), world camera(yellow) and the external arena camera (cyan)) during a freely-moving session. **(f)** Animals wearing CEREBRO exhibited no impairment in their free-moving characteristics such as percent movement, distance covered and speed.

#### Head-Piece Module

The head-piece weighs ∼20g and comprises five different components (*fig 1a*): (1) the scaffold: a curved metal tube that serves as an anchor for all the pieces, (2) eye-cam: an HD MIPI camera (90fps) with Visible light filter and a Macro lens for looking at the eye, (3) world cam: an HD MIPI camera (60fps) for capturing the world scene in front of the camera, (4) an IR LED to illuminate the eye, and (5) a strategically placed Hot mirror to image the eye. The mechanical parts for the head-piece are made up of a Titanium alloy (Ti-6Al-4V) due to strength and weight considerations, and are held to the scaffold using M2 and M4 screw sets (see Supplementary Information 1 for detailed assembly description). To achieve reliable eye-tracking in a fully unrestrained animal, the eye camera’s position requires to be fixed with respect to the animal’s eye. This is accomplished by attaching 2 vertical headposts (5 mm diameter, 1 cm tall cylinders with a flat cut on one side) on the animals’ head (*fig 1c*). The two headposts restrict any rotation or translational movement of the headpiece and allow the headpiece to be placed at the same position for every recording session.

#### Backpack Module

The backpack module weighs 40g and comprises two Customized PCBs, as well as the casing and harness worn by the animals (*fig 1b*). Each camera has its own dedicated PCB (*fig 1d*). Both cameras (eye cam and world cam) have long flex cables (15 cm) and are operated by the PCB.

#### Customized Printed Circuit Boards

The PCB used here is an embedded system that runs on an STM32H750 microprocessor (*fig 1d*). The custom PCB consists of a 0.96” SPI TFT display to preview camera frames and store data image stream and IMU data on a SD card. For more details of the PCB and data storage logic, see Materials & Methods (Printed Circuit Board and data storage).

#### Eye-Tracking System Wearability

When fully configured on a marmoset (fig 1e) the CEREBRO allows a full range of motions characteristic of the marmoset’s natural behavior, while monitoring its position in the environment, eye position, and view of the scene (*fig 1f*). Overall, CEREBRO weighs ∼60 g (headpiece module: 20g + backpack module: 40g), which is comparable to the weight of two infant marmosets that an adult would normally carry on its back. The habituation protocol for adjusting the animals to carrying the backpack and the head-piece is discussed in Materials & Methods (Habituation of animals to backpack). To test the influence of this added weight on mobility, we compared marmosets’ behavior both with and without CEREBRO in multiple sessions (30 min) of freely-moving exploration in a large open arena, (200 cm x 100 cm x 240 cm). The animals were tracked using Optitrack imaging systems (see Methods). Using the velocity threshold of 5 cm/sec, we quantified the percentage of time animals spend moving through the arena. Over 11 sessions spread across months, we observed no significant difference (*fig 1g*) in the time animals spent locomoting vs stationary and scanning the environment (p=0.83). Even though the animals moved for the same amount of time, there is still a possibility that the weight can impede the speed of the animal. Upon investigation, we did not observe any significant difference in either the distance covered by the animals (p=0.569) or the average speed of the animal with and without CEREBRO (p=0.5692). This leads us to conclude that our system does not significantly impede the animals’ movement and can be comfortably used with a freely behaving animal.

### Fast, efficient and accurate pupil detection

In the traditional experimental paradigms that have dominated primate neuroscience, animals are head-fixed and the light source to illuminate the eye can be controlled. The consistency of the light allows for highly accurate eye tracking to be achieved by thresholding the dark iris observed under infra-red light from a brightly illuminated image and the pupil detected either using its centroid or center of the ellipse fit. A critical challenge for eye-tracking in freely moving animals is the lack of control over the illumination. As an individual moves through any real-world environment external lighting conditions and shadows are constantly changing. This point is illustrated by our application of thresholding for pupil detection of a freely-moving marmoset (*Fig 2a*), which can lead to labeling of shadows at the edge of the eye rather than the pupil. Previous studies that examined gaze in free-moving primates relied on conventional pupil thresholding (13–15), which limited the precision that could be obtained. To achieve accurate primate eye-tracking in natural, freely-moving conditions, an alternative method for pupil detection was needed.

**Fig 2:**
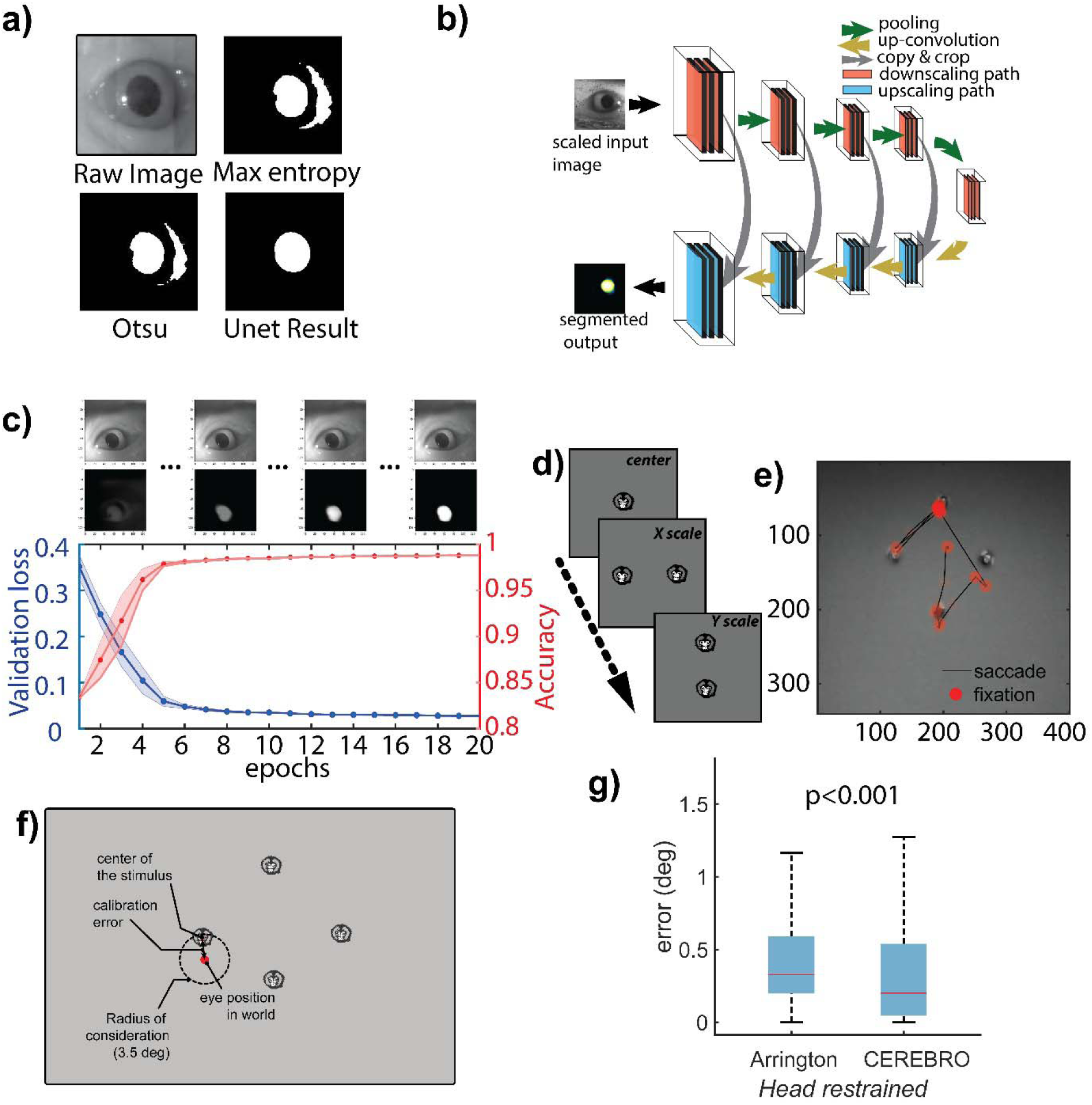
Iris/Pupil detection using segmentation Artificial Neural Network. **(a)** Conventional thresholding approach for pupil detection fails in scenes with variable light intensities. **(b)** Architecture of the segmentation neural network to crop out the iris. **(c)** The performance of the ANN matches closely to that of human annotation with relatively low training epochs (10 in this example) and relatively small amount of training data (∼250 samples) **(d)** For calibrating the eye and world camera, the animals are presented with marmoset faces on a 120 Hz LCD monitor in a predetermined layout. **(e)** The calibrated eye movements can be overlaid on the world scene to determine the visual scene falling on the retina of the animal. (f) An illustration of the error estimation algorithm. **(g)** Accuracy comparison between Arrington eye tracker system and CEREBRO.

Artificial Neural Networks (ANNs) offer an alternative approach to pupil detection under real-world conditions because they rely on features of an image and not necessarily the grayscale values, and therefore are resistant to brightness changes and/or shadows/occlusions. Such approaches have been successfully applied using commercial software, such as DeepLabCut (DLC), to track the pupil of freely-moving mice (16). To further optimize pupil tracking in freely-moving marmosets, here we developed a custom semantic segmentation ANN called UNet (*fig 2b*) (20) that yielded superior performance in these conditions. This workflow allows to detect pupil features robust from lighting and noise (*fig 2b*). With a fully trained network, we reliably detected pupils across various sessions and different animals. The network is easily trainable with fewer epochs for marmoset eyes and a training dataset of only 250-500 images (*fig 2c*). The data preparation, training of the ANN, post-processing, and the use of a user-friendly GUI are explained in detail in *Supplementary information 4* (*Supplementary fig 4b*). This UNet approach closely follows an architecture used previously to segment the pupil from images of human eyes (e.g. RITnet (21). We compared the performance of UNet to the RITnet model (Supplementary Fig. 5). The UNet performed better at identifying the pupil from marmoset eye images, including novel views that varied in position and lighting, but was less robust on human eye images, which is consistent with the respective emphasis in the two models’ training sets. Future modifications of UNet could include additional architecture features of the RITnet model to potentially improve performance.

Accurately calculating the gaze point from the world camera necessitates that the pupil position from the eye camera be calibrated to the real-world position. To this end, we developed the following procedure. The animal is chaired and head-fixed in front of a computer screen placed ∼35 cm directly in front of the subject. Because marmosets naturally look at faces (11), 1-12 small marmoset faces are presented as calibration targets at multiple locations on the display monitor (*fig 2d*). A custom designed graphical user interface was used to adjust for scaling and offset in horizontal and vertical axes by a human operator offline. This allowed us to reliably map pupil eye position to screen coordinates (*fig 2e*), which when head-free, generalized to world coordinates in front of the marmoset. The GUI for pupil detection can be found at https://github.com/Vickey17/UNET_implementation_V2. To compare the system against other pupil-based eye trackers we computed the root-mean-square (RMS) stability of eye position during stable fixation epochs when animals were head-fixed. We find that across 7 recording sessions the RMS stability was 0.05 deg (+/-0.0012 std). These estimates provide a lower bound on the system precision that rivals head-stabilized pupil-based eye trackers such as Arrington eye trackers. To estimate our accuracy with respect to the visual stimuli on the world camera, we created an error estimation algorithm (see Materials & Methods: Error estimation for eye in world calibration) wherein we measure the distance between the calibrated eye position and the visual stimulus (a marmoset face; *fig 2f*). For comparing CEREBRO with the Arrington eye tracker, subjects were head-restrained and presented with marmoset faces on a monitor placed 35 cm from the animal. We observed that CEREBRO had significantly better accuracy as compared to the Arrington eye tracker system (*fig 2g*).

To estimate the accuracy of eye/gaze position in a freely moving animal, we designed a novel paradigm (Fig 3a). Details of the setup are described in the Materials & Methods (Freely moving eye calibration setup). Briefly, the testing arena consists of a plexiglass box with a perch for the animals to sit on. All sides of the box are painted with black acrylic to make it completely opaque except the front panel which is covered with an IR dichroic filter that allows IR light to pass for video recording of the animal inside while blocking the visible light for what the animal can see. A small window (10 x 10 cm) is cut in the filter to allow the animal visual access to see the screen outside. While the CEREBRO equipped animals sit in the dark, a top-mounted galvanometric LASER system draws geometric lines and points on a screen placed ∼75 cm from the box. Each shape is drawn for a duration of 3 sec followed by a dark period of 3 sec. The LASER draws subject’s gaze to these shapes, thereby allowing an estimate of the eye target calibration and its error. In a second set of trials, we also used a single point that moved slowly (<5 visual degrees per second) along random trajectories. In either set of saccade or smooth pursuit trials, the gaze is projected onto the coordinates of the screen through the world view camera and error computed as the nearest distance to a point or line within 3.5 visual degree maximum bound to register looking at the feature (*fig 3b*). Projected eye position was found to track geometric shapes defined by points and lines (*top panels, fig 3c*) and during pursuit of the point (*bottom panels, fig 3c*). To compare accuracy against the head-restrained preparation we presented the same stimuli to head-fixed marmosets. We observe that errors cluster below 2 visual degrees in all conditions, well away the maximum bound for registering looking at a feature, and the median accuracy remained less than 1 degree in both saccade trials for geometric shapes (*fig 3d*) and smooth pursuit trials for a moving point (*fig 3e*). The median error was higher for both trial types in the freely-moving compared to head-restrained condition with an increase in error by 1.98 times for saccade trials and 1.4 times for pursuit trials (Wilcoxon rank sum test, p<0.0001 saccade; p=0.006 smooth pursuit), but still remained under 1 visual degree, validating an upper bound of the system’s accuracy.

**Fig 3.:**
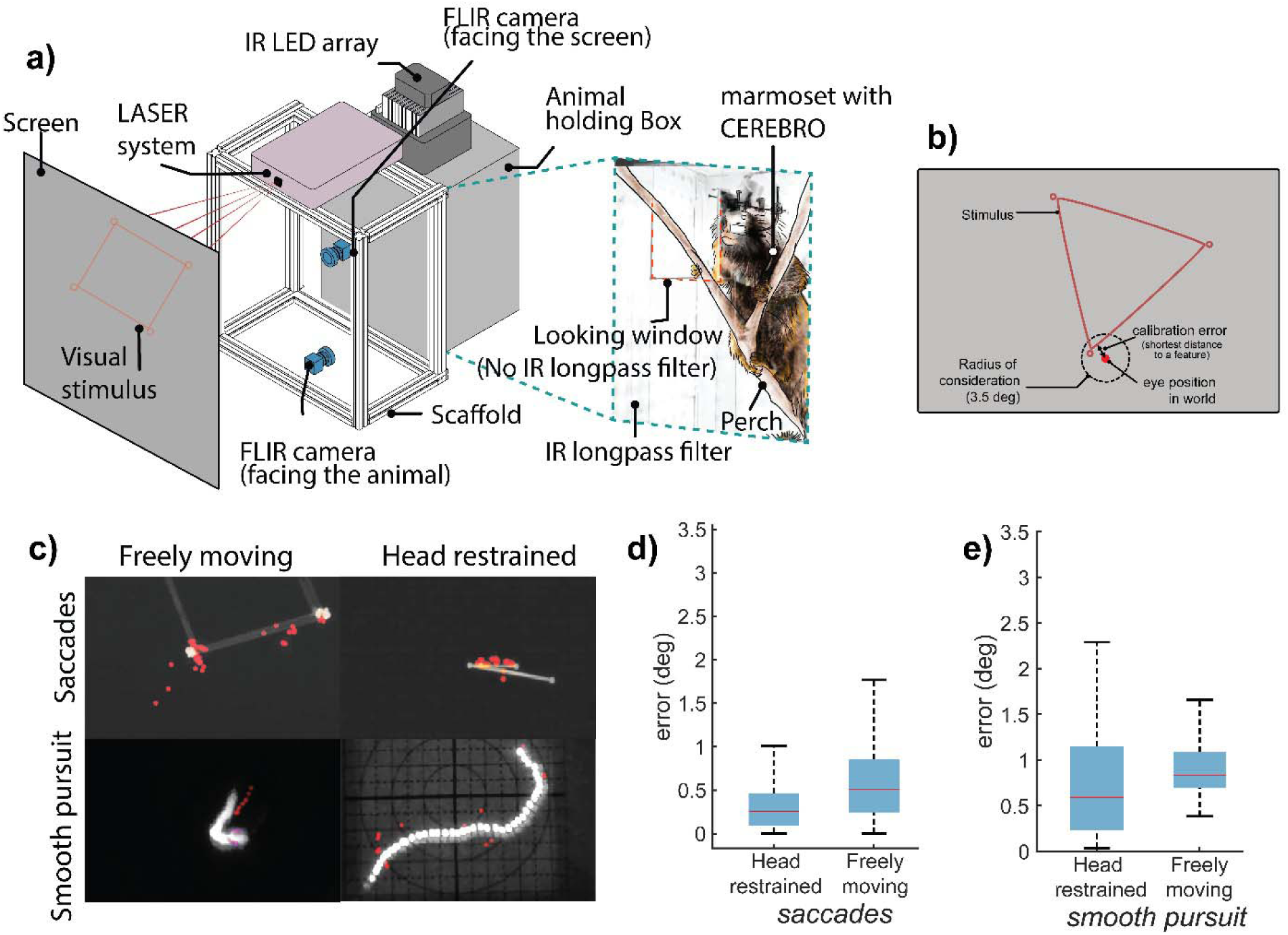
Eye/gaze accuracy in various experimental setups using CEREBRO. **(a)** Schematic for the experimental setup used to test accuracy of eye calibration in unrestrained animal. The setup consists of a dark box with a perch. The front of the box is covered with an IR longapss filter that doesn’t allow visible light to pass except for a small window shown with red dashed line in the inset image. A galvanometric LASER system draws different geometric shapes on a screen for the animal to observe. **(b)** Schematic illustrating error estimation algorithm for freely moving eye calibration. **(c)** Maximum intensity projection of stimulus and eye in world position (red markers) for saccades and smooth pursuit trials in freely moving and head-restrained setup. **(d)** Accuracy comparison for eye calibration error for head restrained and freely moving animals during saccade trials and **(e)** smooth pursuit trails.

### Electrophysiology in combination with CEREBRO

CEREBRO was designed explicitly to create a tool to investigate the neurobiology of natural, active vision in freely-moving monkeys. As such, integral to its design was the capacity to simultaneously record neural activity, eye behavior, and the visual scene of the animal, as each are integral to this broader ambition. We tested the validity and sufficiency of the system to this end by performing experiments with CEREBRO while activity of single neurons was recorded with chronically implanted multi-electrode arrays (N-form, Modular Bionics) in the primary visual cortex (V1) of the marmosets. These experiments sought to (a) estimate visual tuning properties of neurons in primate visual cortex, i.e. receptive field mapping, as well as orientation and spatial frequency tuning (22–25), using CEREBRO in more traditional head-fixed paradigms, so as to demonstrate the accuracy of our eye-tracking system by replicating these classic effects, and (b) obtain eye, head and body behavior, visual scene, and activity of single neurons simultaneously in freely-moving paradigm to demonstrate the capacity of CEREBRO in investigating primate active vision.

To recapitulate the tuning properties of V1 neurons, subjects were head-fixed while wearing CEREBRO and presented with the following stimuli: flashing dots for receptive field mapping, and drifting gratings with different orientations and spatial frequencies for tuning properties (*fig. 4a*; see online methods). Critically, marmosets were allowed to free-view the video screen during stimulus presentation and offline corrections for eye position enable accurate reconstruction of visual properties following a recently developed free-viewing approach (26). A key difference here from the previous study with head-fixed marmosets is that the visual input is obtained by the world camera’s view along with the estimated eye position from CEREBRO instead of what was known to be displayed on the screen, thus validating that these methods could generalize to real-world stimuli. Results from three example neurons demonstrate visual receptive fields estimated at the peak visual latency (*fig. 4b top row*, Online Methods). The orientation tuning at the peak visual latency (*fig. 4b bottom row*, Online Methods) estimated from the CEREBRO world camera image. These results replicate classic findings about the properties of neurons in early visual cortex thereby demonstrating that our eye-tracking system and calibration approach can accurately record neural responses in response to visual stimuli on the primate retina.

**Fig 4:**
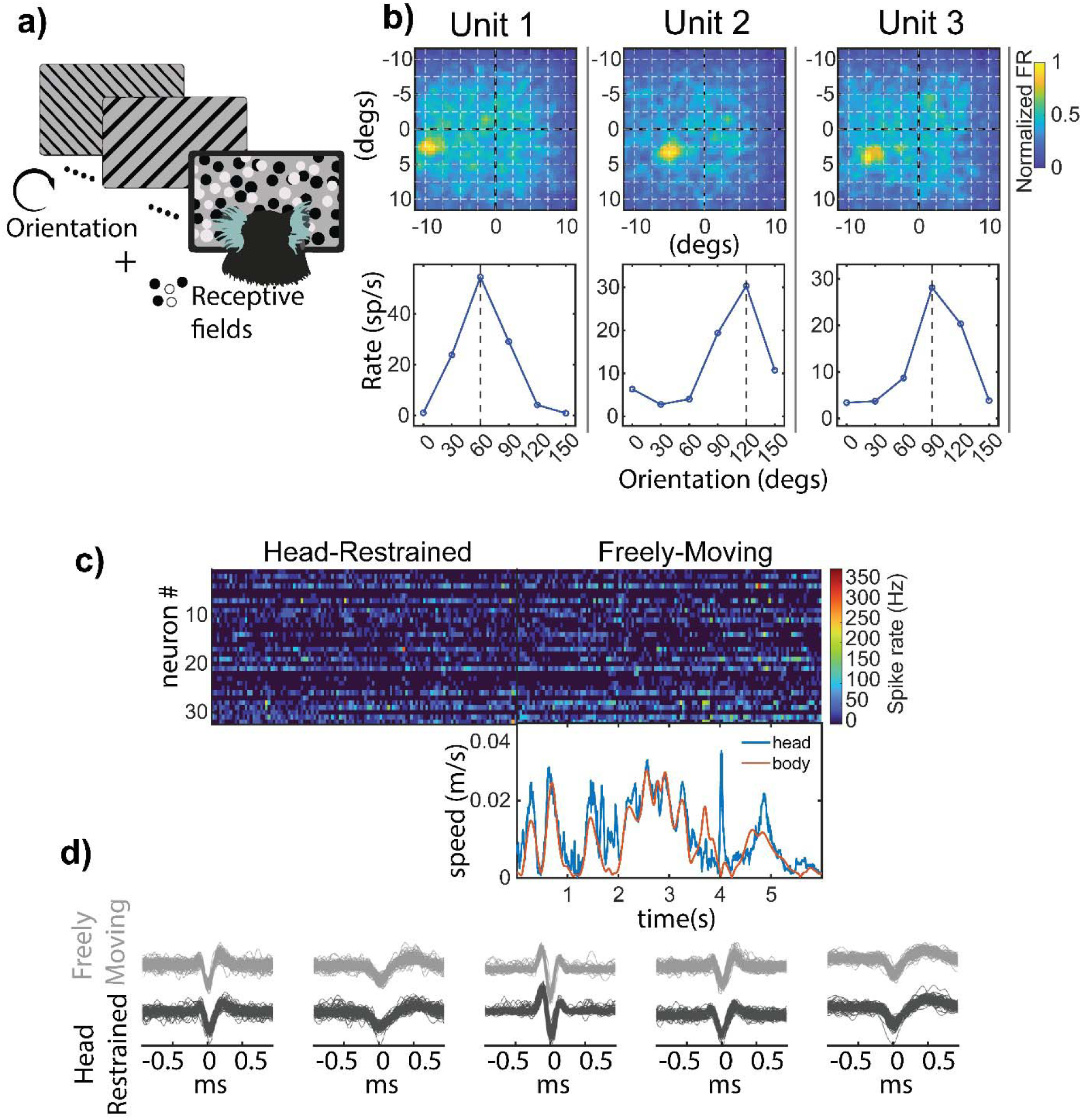
Eye tracking coupled electrophysiology in freely moving marmosets. **(a)** To validate use of CEREBRO with electrophysiology, subjects were presented with orientation grating and receptive field stimuli in a classical head fixed preparation. **(b)** The receptive field (top) and orientation tuning curve (bottom) is shown for three representative neurons recorded in marmoset V1 while using CEREBRO. **(c)** Example raster of 30+ single neurons in marmoset V1 while subjects are head-restrained (left) and freely-moving (right) while wearing CEREBRO. The head (blue) and body (purple) speed of the marmoset using CEREBRO is plotted below the freely-moving raster **(d)** Example spike waveforms from five exemplar neurons demonstrate that we were able to stably record from same neurons in head-restraint vs freely moving conditions.

Following this, the animal was allowed to actively explore a 200cm x 100 cm arena decorated with various visual stimuli. Activity of single neurons was continuously recorded in the period of head-restraint and freely-moving condition; body and head movements were simultaneously recorded using *OptiTrack* system (*fig. 4c*, Online Methods). The comparison of spike waveforms in head-restraint versus freely-moving condition among 5 example neurons demonstrates the stability of neural recording throughout the session (*fig. 4d*). Within the same session, the eye behavior and the neural activity (spike rate) change between the head restrained vs freely moving animal. We explore some of these behaviors in the following sections.

### Visual behavior of freely moving marmosets

We compared the eye behavior of a head-restrained marmoset with that of a freely moving marmoset. In the head-restrained context, the animal was presented with a series of naturalistic images (see Materials & Methods). During the freely-moving context, the animal was placed in an open arena and allowed to explore the environment. Consistent with previous studies (11), analysis in the head-restrained context revealed that the eye in head position i.e. the horizontal and vertical eye position spanned +/-5 visual degrees for 2SD of all the positions (fig 5a) and the eye position distribution for saccades vs fixation were identical in this case (fig 5b & c). However, the results for freely-moving marmoset are starkly different. Firstly, the eye in head position is much more limited in a freely moving animal spanning only +/-2.5 visual degrees (fig 5d). Moreover, most of the eye positions at the edges of the range were involved in gaze shifts and not gaze fixations (fig 5e & f). These findings suggest that during stable epochs of gaze the head-position provides a reliable estimate of gaze position. The primary motivation to develop CEREBRO was to precisely quantify the characteristics of eye-movements and gaze in freely-moving, naturally behaving primates. To this end, we recorded visual behavior in marmosets wearing CEREBRO as they explored an open rectangular arena (*fig 6a*). As marmosets do not continuously move when in open-field environments, we distinguished between the following two behavioral states in these test sessions to determine whether differences in visual behavior emerged: a) “*stationary”*-the monkey was seated or standing and visually scanning the environment without physically changing locations (b) “*locomotion*”-the monkey was physically moving and changing its position in the environment (Online Methods). In a typical session, a marmoset remains stationary for extended periods at fixed locations (*occupancy map in fig 6a*), and then moved between those locations (*gray traces, fig 6a*). We conjectured that the gaze dynamics likely differs between these two behavioral states, as each differs in motoric demands and exploratory function.

**Fig 5.:**
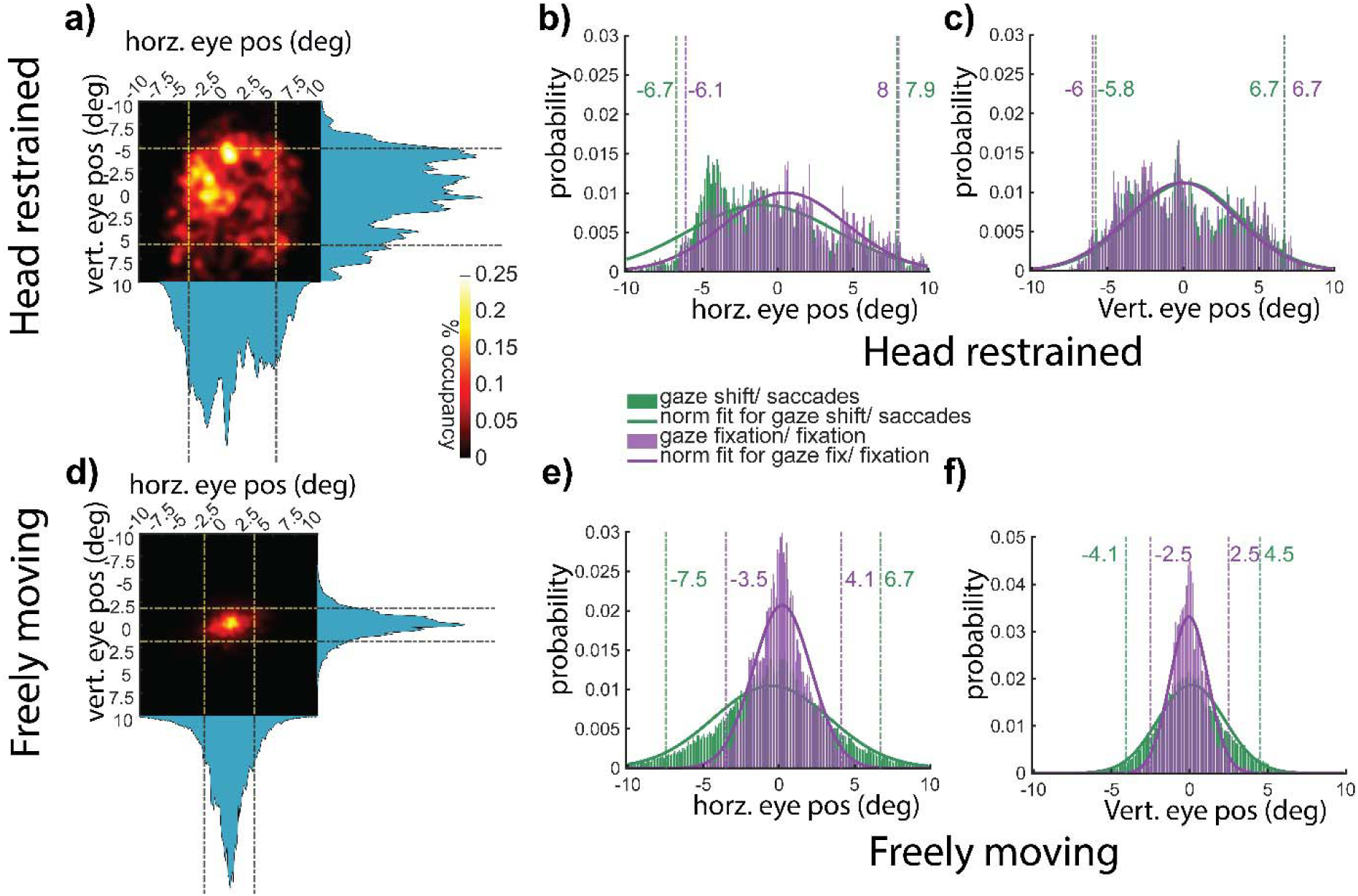
Characteristics of eye/gaze behavior in various experimental setups using CEREBRO. **(a)** Heatmap of eye position in a restrained marmoset with dashed lines marking 2 SD mark. **(b & c)** distribution of horizontal and vertical eye position split by fixation (purple) vs saccade (green) with corresponding dashed lines representing 2-SD of data variation. **(d)** Heatmap of eye position in an unrestrained marmoset. **(e & f)** distribution of horizontal and vertical eye position split by gaze fixation(purple) and gaze shifts (green). Color-matched dashed lines represent 2-SD of variation in data.

**Fig 6.:**
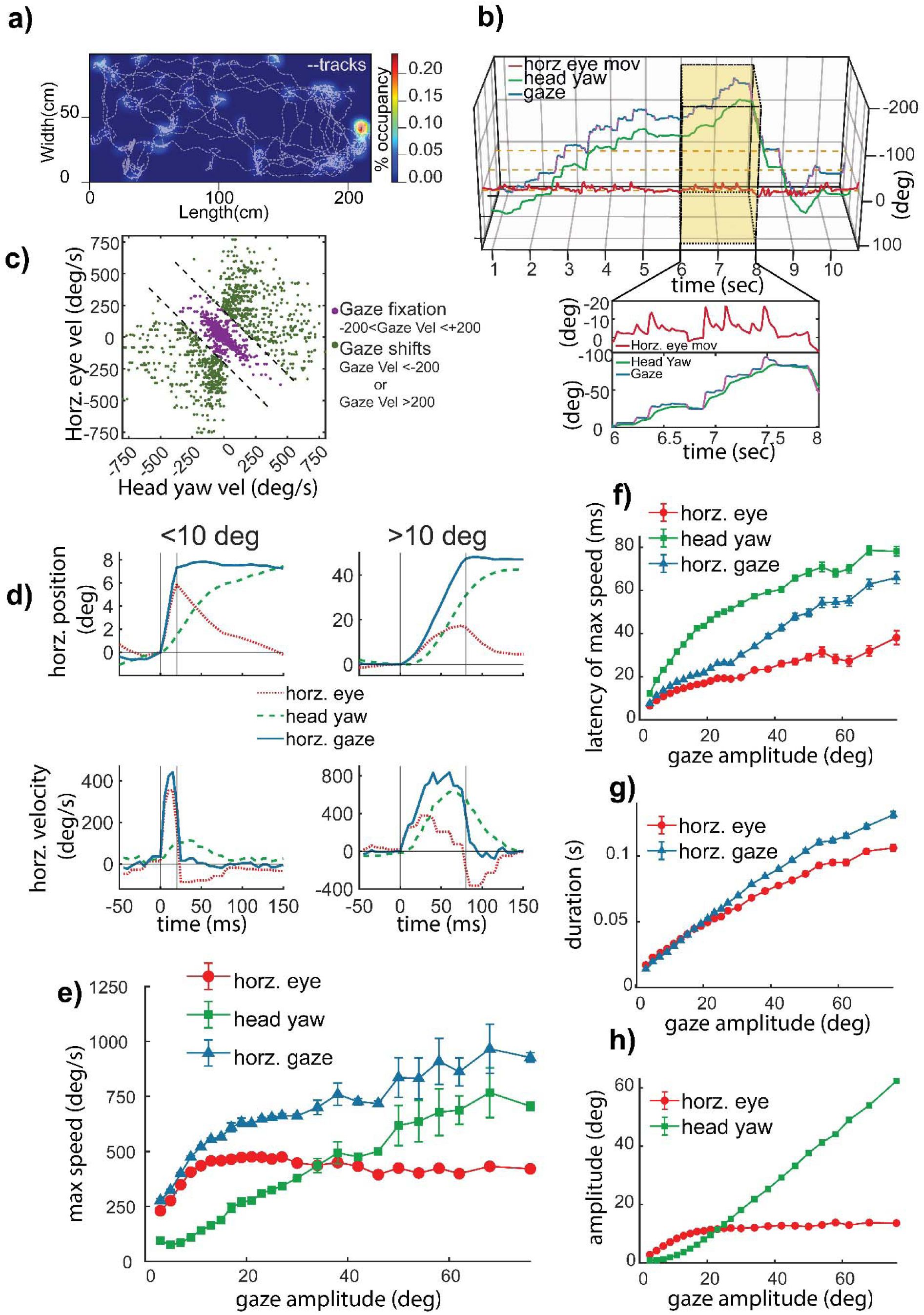
Gaze characteristics in freely moving marmosets. **(a)** Representative session of activity pattern for marmosets in an open arena (200 cm x 100 cm x 240 cm). Grey dotted lines indicate periods of locomotion, while heat maps plot stationary occupancy. **(b)** Plots the estimate of the animal’s gaze (blue) with CEREBRO. Horizontal eye movement (red) and head yaw (green) are also plotted. Inset magnifies a 2s period of time. **(c)** Horizontal eye movement and head yaw in freely-moving marmosets are shown. Green dots indicate gaze fixations, while purple dots indicate gaze shifts. **(d-h):** eye-movements (red), head-movements (green) and gaze-shifts (red) in freely-moving marmosets. **(d)** Plots the position (top) and velocity (bottom) of eye-movements, head-movements and gaze-shifts <10degrees (left) and >10degrees (right) **(e)** Shows the maximum speed and gaze amplitude. Plots the latency to max speed (f), duration (g) and amplitude (h) as a function of gaze amplitude

In a freely moving primate, visual exploration is accomplished by changing gaze which can be defined as a sum of head and eye movements (*fig 6b*). Stable eye positions in this context are rare because even when an animal fixates at a fixed position in the scene – referred to here as a gaze-fixation-the eyes must still move smoothly to compensate for any head-movements for retinal stabilization. These compensatory eye movements reflect the vestibular ocular reflex (VOR) that well conserved across species and normally engaged to reduce retinal motion that is due to head and body motion (8, 9, 27–29). These compensatory movements correlate negatively with the head-movement velocity to subtract its effect and achieve stable gaze. By contrast, during rapid gaze-shifts, equivalent to traditional saccades in the head-fixed case, the VOR is suppressed and there is a combination of conjugate head and eye movements along the same direction, with eye velocity reversing at the end of the rapid shift as VOR is restored and gaze is again stabilized by compensatory eye movements compensating for continuing head velocity. In our current study, we have primarily focused on the horizontal eye movements and head yaw since it has been demonstrated that horizontal pursuit eye movements are more accurate and symmetric than vertical ones in primates (30, 31). The inset in Figure 6b illustrates this point as the sum of eye and head position (gaze) exhibits steps in position with stable periods in between. While head position is continuously changing, the eye position exhibits saw-toothed type patterns in which a jump in position is followed by decay backwards that compensates for the change in head position. To distinguish epochs of rapid gaze shifts and compensatory movements, we set a gaze velocity threshold of ±200 °/sec. Examination of eye movements during gaze-shifts and gaze-fixations revealed a clear trend of negative correlation between head and eye movements for fixation periods reflecting compensatory eye movements to stabilize gaze (*purple, fig 6c*) in a freely-moving marmoset. For the gaze velocity threshold set, we find compensatory movements are well separated from other rapid gaze shifts (*green, fig 6c*).

A core feature of mammalian ocular-motor behavior is known as the main sequence; a characteristic linear relation between the amplitude and peak velocity of eye movements. To test whether the main sequence is evident in a freely-moving primate, and characterize it with respect to head and gaze, we quantified the main sequence and amplitude-duration relationship. For conjugate gaze shifts we observed a characteristic pattern wherein eye velocity reached a peak velocity more quickly than head-velocity. As head-velocity decayed with a long tail, eye velocity reversed direction in order to counter-act the head-velocity and stabilize gaze (*fig 6d*). This pattern was evident both for small and large gaze shifts, with the duration of the gaze shift being longer for larger gaze shifts. To quantify these patterns more accurately, we next plotted the peak velocity, latency to peak, and duration of the eye, head, and gaze components as a function of gaze amplitude (*fig 6e-h*). Although the main sequence was evident in freely-moving marmosets here, evidence suggested differences in how the eye and head components contribute to it as a function of gaze amplitude (*fig 6e*). Whereas the peak velocity of eye movements saturated for gaze shifts of roughly 10 degrees in amplitude around 400 degrees/sec, the peak velocity of head shifts continued to increase even for the largest measured shifts out to 80 degrees. The latency of the peak eye velocity always leads peak head velocity as a function of gaze amplitude with each growing as a function of shift amplitude (*fig. 6f*). The peak of the gaze velocity follows more closely with eye velocity for small shifts (<36 degrees) and with head-velocity for larger shifts. The duration of gaze shifts follows a roughly linear relation with gaze amplitude (*fig. 6g*). In summary, the components of eye and head shifts differ in their contribution to gaze shifts depending on the amplitude of the shift, with eye velocity rapidly saturating in its maximum velocity for shifts of about 10 degrees in size after which the slower initiating head-shift contributes more to the total gaze movement. As the gaze amplitude becomes larger, the contributions of the head increase linearly to the overall shifts while the eye saturates in its contribution after about 20 visual degrees (*fig 6h*).

To determine how visual behavior differed between behavioral contexts we compared the eye-movements of marmosets when head-fixed and freely-moving, distinguishing between instances when individuals were visually scanning the environment while ‘stationary’ and instances when animals were actively moving through the world by ‘locomotion’ in the latter context (*fig 7a*). As demonstrated in *fig 7b*, differences in marmosets’ visual behavior when head-fixed and both freely-moving contexts were stark, with the eye movements being notably dynamic in freely-moving contexts reflecting VOR adjustments for self-motion. We quantified these differences further by calculating the approximate entropy of marmoset eye movements in each of these three contexts (32). This analysis estimates the randomness of a time series where higher value of approximate entropy means a system that is more random and vice versa – thereby allowing a metric for quantification of the visual behavior as a time series. Overall, we observed that marmoset eye-movements in the head-restrained condition exhibited a significantly lower entropy than both the freely-moving conditions (stationary and locomotion; *fig 7c*), suggesting that eye movements in a freely moving animal is more chaotic/random. However, the approximate entropy for “gaze” of freely-moving animals – the combination of both head and eye-movements – a different pattern emerged. Here the approximate entropy in both freely-moving contexts was in fact lower than in the head-fixed context. This suggests that the synergistic combination of head and eye movements in a freely-moving animal yields a less random/chaotic scanning of the visual environment than when head-fixed, a likely computational optimization of the visual system to sample the scene in animals as they naturally move and explore the world populated with elements with varying salience. The statistics of maximal speed and amplitude of the eye movement shows a significant faster and larger eye movement in the freely-moving context (*fig 7d*).

**Fig 7.:**
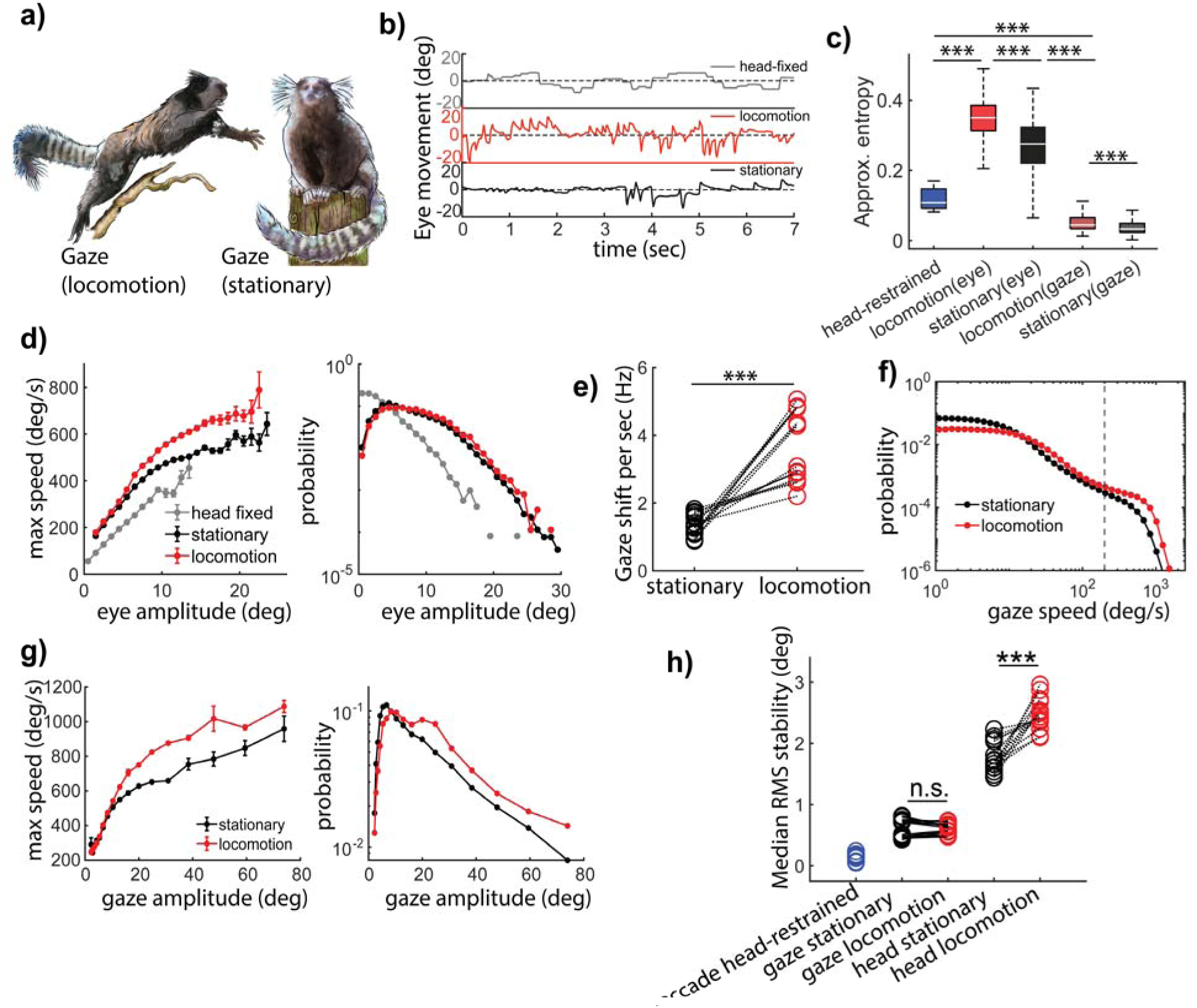
Characteristics of horizontal eye movements in freely moving marmosets. **(a)** An illustration of the gaze shifts in a freely behaving marmoset in two different states of locomotion(left) and stationary(right). (b) Animals exhibited noticeably different eye behavior in various contexts i.e. chaired, locomotion and stationary. **(c)** Horz. eye movements have significantly less “approximate entropy” in head-fixed context as compared to freely moving. Although horz. gaze has the least approximate entropy of all the contexts. This shows that a freely moving gaze is computationally more regular than a head-fixed context. (d) Freely moving animal exhibits larger range of horz. eye amplitude and the speed of horz. eye movements is higher during locomotion than stationary state. Marmosets make more gaze shifts (e) during locomotion with higher speed (f) and larger amplitudes (g). (h) RMS stabilization of head(yaw) and horz. gaze shows that despite poorer stability because of head movements, gaze remains remarkably stable during locomotion and stationary epochs.

In addition to significant contrasts in visual behavior between head-fixed and freely-moving contexts, we also observed differences in the latter when marmosets were stationary or locomoting. Specifically, marmosets make significantly more gaze shifts per second (*fig 7e*) with greater speeds (*fig 7f*) and larger amplitudes and maximal speed (*fig 7g*) during locomotion than during stationary phases. This pattern raises the question if the gaze of the animal is less stable during locomotion given the larger gaze shift amplitudes. To investigate that, we calculated the Root-mean-square (RMS) stabilization of the gaze during gaze fixations during locomotion and stationary contexts (16). We use RMS as a measure to quantify the deviation of gaze and head yaw during stabilization periods (gaze fixations). A smaller RMS during fixation epochs indicates a more stable fixation while a higher value for RMS will indicate poor stabilization. Analyses revealed that gaze was on average about 3 times more stable than head movements during gaze fixations (*fig 7h*). When comparing the stability of gaze to that of head, however, we observed that although head movements were less stable during locomotion gaze stability surprisingly improved during locomotion (*fig 7h*), despite the increase in head-motion. Thus, while the head is clearly less stable during locomotion, compensatory eye movements appear to provide better stabilization achieving more stable epochs of gaze fixation than even the sedentary phase. This highlights the importance and potential context-dependence of gaze stabilization, wherein it appears to be enhanced to achieve greater stability during locomotion.

## Discussion

Here we quantified the active visual behavior of a freely-moving primate by developing an innovative head-mounted eye-tracking system. Our system achieves several technical innovations that enabled accurate quantification of head and eye movements in a small bodied (∼300-400g) monkey – common marmosets-with the resolution and speed needed to accurately quantify primate visual behavior in real-world contexts. Specifically, this system achieves high-speed recording of the eye (90 FPS) and world (63 FPS), as well as applies an innovative solution to overcoming the challenge that frequent changes in lighting cause for pupil detection by leveraging a segmentation neural network. As CEREBRO was designed to seamlessly integrate with wireless neural recording methods, this system is poised to not only elucidate the dynamics of active visual behaviors but the supporting neural processes. While findings reported here in freely-moving primates recapitulate the core features of conjugate gaze movements from prior studies using head-free-but chair-restrained – macaques (33, 34) and extend those to freely moving primates, analyses also revealed several related characteristics of visual behavior that have not been observed previously. Perhaps most notably, we observed that the entropy of the gaze behavior was significantly lower when animals were freely moving than when head fixed, and that the stability of the visual gaze improved during locomotion. Overall CEREBRO opens the door to examining active primate visual processing in the same naturalistic, freely-moving contexts in which the system has been optimized over evolution, wherein coordinated head-eye movements act to stabilize vision during natural exploration, critical features that cannot be recapitulated in the absence of ethological movements.

CEREBRO is designed to allow for studying visual behavior coupled with neural responses in a freely moving animal. This will enable investigation of the neural correlates of active vision. To that end, we tested our system during neural recordings to characterize orientation tuning and receptive fields using classical reverse correlation methods during head-fixation. As demonstrated in Fig. 4, we were able to recover clear receptive fields and orientation tuning of V1/V2 neurons using eye tracking from CEREBRO in combination with the view from the world camera. In the future, this system will more broadly enable us to investigate the role of saccade related modulation in head-fixed vs head-free contexts, and more so, to obtain receptive fields in freely moving animals. Further, the video streams from our world camera will provide statistics for how the visual scene and optic flow changes during free motion and help to expand our understanding of the natural scene statistics and how they are processed in vision.

More generally, it will be possible to estimate visual receptive field properties up to the accuracy of the gaze tracking in freely moving scenarios. Our estimates of gaze accuracy, though less accurate in the free-moving than head-fixed conditions (fig 3d and 3e), are still roughly comparable and on median under 1 visual degree. Thus, reconstruction of visual receptive fields, as was shown for the head-fixed condition using the scene camera (fig 4b), should also be feasible for free-moving cases given similar flashed visual stimuli. However, a key challenge during free-motion is that the statistics of the visual input are radically different and instead controlled by the animal, which on one hand provides a novel and exciting aspect of active vision but on the other hand pose challenges for fitting receptive field models due to the spatiotemporal correlations found during natural motion and with real visual scenes. These issues can be addressed by exploiting computational approaches that are highly data-efficient. Indeed, the receptive field mapping of V1 neurons in freely moving mice, where visual receptive fields are much larger than primates, has been achieved with a Generalized Linear Model (GLM) approach (3). Similar methods could be applied in free-moving marmosets, especially if the visual areas under study have receptive field sizes larger than gaze tracking error, as in the peripheral visual field.

The contribution of eye and head movements towards conjugate gaze shifts in freely-moving marmosets were qualitatively similar that of chair restrained but head-free macaques and other mammals (7, 33, 35–39). Although broadly similar, marmosets did exhibit some quantitative differences relative to macaques, potentially due to their smaller head-size. For example, marmoset eye movement velocity and amplitude saturate for much smaller gaze shifts around 10-20 degrees, after which the head contributes to the bulk of the shift in gaze (Fig. 4h), whereas in the macaque this transition does not occur until one reaches much larger gaze shifts between 20-40 degrees (33). Moreover, most gaze shifts in macaques under 20 degrees in size are predominantly driven by shifts in eye position and not the head, while in the marmoset the same gaze shifts would have significant head-movement components. Only the smallest of gaze shifts under 5-10 degrees in the marmoset, a range comparable to the movements made in head-fixed marmosets (*fig 6d*), are dominated by changes in eye position after which the head would normally make significant contributions. These differences likely reflect an efficiency tradeoff related to smaller head-size in marmosets (40).

Because we recorded marmoset eye-movements in head-fixed and freely-moving conditions, this novel dataset can both be compared to prior studies and extend our understanding of visual behavior in this New World primate. As in previous experiments (11), when head-fixed the oculomotor range is relatively fixed within about 10 visual degrees. But when marmoset is freely-moving that range increases to roughly 20 degrees (*fig. 7d*), suggesting the limited oculomotor range when head-fixed represents more of a motor preference than a physical limitation. In a previous study of chaired but head-free marmosets, a paradigm was used to evoke large gaze shifts involving up to 180-degree head rotation (12). Although similar large shifts were rare in the study here, a similar linear relation between peak head velocity and gaze shifts that peak near a velocity of 750 °/sec for an 80-degree gaze shifts were evident in freely-moving marmosets suggesting that this aspect of head-gaze control are relatively invariant to the form of the task. A direct comparison of the two freely-moving contexts here revealed a number of notable differences including larger amplitude gaze shifts with higher maximum velocities when locomoting (*fig. 7, d-g*). By contrast, when stationary and visually scanning the environment, marmoset biased to smaller gaze shifts indicating that motor demands of actively moving through space likely drives differences in how the head and eyes coordinate to stabilize the visual field.

Our understanding of NHP vision is almost entirely based on studies in which subjects are head-restrained. Quantifying primate visual behavior with the innovative eye-tracking system here afforded the powerful opportunity to examine whether our assumptions about natural vision are accurate. Indeed, we observed broad qualitative similarities in many features of conjugate eye movements between head-fixed and freely-moving contexts. However, it was also apparent that certain assumptions about results in the more conventional paradigm may not be strictly true. The first being that gaze stability was not worsened despite greater eye and head movements during locomotion, and in fact, was slightly enhanced. We observed that the approximate entropy of the marmoset visual system was significantly lower (i.e. improved) when individuals were freely-moving than head fixed suggesting that the collective visual system is more consistent and predictable when marmosets are moving and exploring the environment. Likewise, a comparison of freely-moving marmosets when stationary – i.e. visually scanning-and locomoting revealed that the stability of visual gaze-as measured by RMS – was significantly better when animals were locomoting. In other words, despite an increase in the number, speed and length of the gaze shifts when locomoting, vision actually became more stable. These findings show that the coordinated movements of head and eyes have been optimized to accommodate self-motion in a diversity of ethological contexts.

The context-dependence of visual stability between the stationary scanning and locomotion states could reflect changes in VOR at the neural level, as well as other positional strategies that optimize the VOR system (41, 42). Such context-dependence of gaze control has long been appreciated from head-mounted eye tracking studies in humans (43, 44), and recently includes free motion in natural contexts (45). The technical parallels between these eye-tracking systems in human experiments and CEREBRO allows for comparisons between these primate species, as well as between marmosets and other taxa. In humans, for example, the gain for angular VOR (aVOR) is a function of locomotion speed(46, 47). Those findings are consistent with the RMS stabilization of gaze in marmosets being maintained, or even slightly improved, during locomotion as compared to when they are stationary and scanning the scene (Fig. 7h). An increase in VOR gain during locomotion would explain how stability is maintained even though head and gaze movements increase in frequency and amplitude during locomotion (Fig. 7e,f,g). Further, we measured the negative correlation between horizontal head and eye velocity during periods of stable fixation to estimate the VOR gain directly (Supplementary Fig. 8). We find that VOR gain increases during locomotion as compared to stationary scanning, consistent with prior human studies and the maintenance of RMS stability during locomotion.

Another parallel across humans and marmosets is the change in the relative contribution of head and eye movements to gaze shifts with increasing gaze amplitude(48). We show that larger head amplitudes lead to less eye contribution (Fig 6h) wherein the eye amplitude saturates at ∼10 deg and then the putative target is achieved with help of head-movement. In humans the reliance on head-movements does not typically occur until much larger gaze shifts are required, but this also can depend critically on the task context, including the speed of locomotion(46). The natural movement statistics of head-position in free-moving primates, however, is known to differ substantially from that of rodents (49), suggesting that at least some mechanisms for visual stabilizations may not be evident across all vertebrates or mammals. In future work the system could be expanded to address other features of oculomotor control that could be especially relevant to free-motion in these contexts in primates and other taxa, including vergence and torsional eye movements (50, 51). The current eye tracking system offers an opportunity to study how computational constraints and movement strategies during active vision act to achieve stability and provide high acuity vision in different contexts, including locomotion and navigation through the environment.

From the outset, CEREBRO was explicitly designed to allow for studying visual behavior coupled with neural responses in freely moving animals. This enables us to investigate the neural correlates of active vision. To that end, we tested our system on classical visual correlates like orientation tuning and receptive fields to assess the quality of responses collected by our system with the conventional methods. As demonstrated in our findings (fig 4), we were able to obtain clear receptive fields and orientation tuning of V1/V2 neurons using just the eye and the world camera of our system. However, quantifying the visual information at the level of the retina and its representation in early visual cortex presents particular analytic challenges beyond the resolution of the eye-tracking system. In contrast to head-fixed paradigms in which controlled visual stimuli can be presented to map receptive fields in visual cortex, the natural environment has highly correlated spatiotemporal features in the visual input, as well as low contrast in a large number of images (e.g. when the animal looks at the ceiling or the floor). These issues cause insufficient statistical power in the collected samples and require more advanced computational approaches to resolve that were beyond the scope of this study. Fortunately, a recent elegant experiment that addressed this issue in mice (3) provides a roadmap that we will pursue in future studies Additional fundamental questions will also be pursued in future experiments such as dissecting the role of visual flow on the neural population to understand how optical flow aids in separating foreground and background elements as previously studied in some of the human experiments (52). Indeed, the innovative system described here eliminates a critical bottleneck for the field that opens the door to studies of natural active vision in primates that may expand our understanding of the underlying neural mechanisms that support our preeminent sensory domain.

Despite its potential, the innovative eye-tracking system described here is not without its limitations. For example, like most primates, marmosets are arboreal. This 3D environment presents significant but distinct challenges to the visual system compared to the 2D environment tested here. However, it is not clear how this wearable technology may affect their movement in more precarious environments, and by extension the data and our interpretations of those data. Moreover, one of the key plans for future research is to leverage this system to investigate social perception and its underlying neural mechanisms (e.g. face patches) but it is not yet clear if the hardware on the head would affect how other marmosets perceive and interact with a test subject, which could have similar biases on the resultant data. Finally, we have made significant efforts to quantify the precision of the marmoset gaze targets in a freely moving context, but it is important to recognize that these are estimations. Although the same can be said of all eye-tracking systems, the inherent complexity and variability of real-world environments have the potential to introduce more error in our quantification than in more controlled experimental settings.

The sensory and motor systems of animals co-evolved. Amongst the most significant selective forces acting on animals is to move and interact with the world, and to do so requires sensory feedback. This ground truth inherently couples sensory and motor systems. The animals’ behavior can be thought of as an outcome filtered by the environment and the animals’ potential actions (53). For a freely moving animal this “affordance landscape” can be very dynamic given the exteroceptive (sensory feedback), proprioceptive and interoceptive feedback. These parallel and complementary feedback processes work in synergy to guide relevant behavior of the animal such as decision-making. It has been argued that perception is not just about creating an internal representation of the world based on sensory inputs, but instead filters responses relevant to the environment and animals’ internal state (54). Despite this, the study of the primate visual system has largely ignored considerations of movement and relationships between the sensory inputs and affordances. Head-restrained experiments limit the sensorimotor and “affordance landscape” of an animal that potentially bias results in a way that differs from naturalistic behaviors in which an animal acts as an agent. Freely moving animals express continuous behaviors based on the elements in the environment, history of previous actions and internal state which is starkly different from a trial-based structure and lead to many interesting discoveries about computation strategies in the brain. Leveraging our innovative head-mounted eye-tracking system for marmosets, we provide compelling evidence that the coordinated actions of eye and head movements are optimized to stabilize primate vision and increase its predictability. These patterns emphasize the significance in considering the ethological relevance of how primates are tested in vision studies. Evidence suggests that neural responses in head-fixed paradigms are not necessarily predictive of how the same single neurons – or population ensembles – respond in naturalistic contexts (55–57). Indeed, investigations of vision in freely-moving mice suggest that at least some elements of neural activity in this context are distinct (3, 58). Our innovative eye-tracking system can be leveraged in marmosets to precisely examine the primate visual system during natural, freely-moving behaviors (5, 59) and address a suite of foundational questions that we have as of yet been unable to investigate with sufficient quantitative rigor.

## Acknowledgements

We thank Alex Huk and Cris Niell for their invaluable insights throughout the development of this system. We thank Brian Corneil for his valuable comments on an earlier version of the manuscript. This work was supported by grants from the NIH/BRAIN Initiative (R01 NS118457 & U01 NS116377) and AFOSR (FA9550-19-1-0357) to CTM and by a Kavli Institute for Brain and Mind Postdoctoral Award to JL.

## Author contributions

Conceptualization: VPS, JL, JM, and CM; Methodology: VPS; Software: VPS; Formal analysis: VPS and JL; Investigation: VPS and JL; Resources: VPS, JL, CM; Writing: VPS, JL, JM, CM; Visualization: VPS, JL, JM, CM; Supervision: JM, CM; Funding acquisition: CM

## RESOURCE AVAILABILITY

### Lead Contact

Further information and requests for resources and reagents should be directed to and will be fulfilled by the lead contact, Cory Miller.

### Materials availability

The STL designs for the headpiece and its material description can be downloaded at https://github.com/Vickey17/CEREBRO_HeadPiece.

### Data and code availability

- Head, eye and body locomotion data has been deposited at https://doi.org/10.5061/dryad.8gtht76xb
- The GUI for UNet segmentation-based pupil detection can be found at https://github.com/Vickey17/UNET_implementation_V2.

## Materials & Methods

### 1. Experimental model and study participant details

The development and testing of CEREBRO were performed on two common marmosets (monkey M, male; monkey S, female). Both subjects were group housed in home cages (76 cm x 76 cm x 81 cm) and were 2 years old at the time of implant. Monkey M had a chronic implant in left V1 and monkey S had bilateral chronic implants in both left and right V1. All surgeries and experiments were performed in the Cortical Systems and Behavior Laboratory at University of California, San Diego (UCSD), and approved by the UCSD Institutional Animal Care and Use Committee in accordance with National Institute of Health standards for care and use of laboratory animals.

### 2. Fitting the animal with CEREBRO

The head assembly for CEREBRO is custom fitted to every animal. For the initial fitting, the animal is anesthetized using a combination of Ketamine (dose:20 mg/kg) and Acepromazine (dose: 0.75 mg/kg). All the parts (*Supplementary fig 1b*) are adjusted for best view of the eye and the front camera and fastened using set screws.

### 3. Habituation of animals to backpack

Subjects were habituated to carry backpacks for 1-3 months. This was done in habituation sessions by placing the backpack on an individual and incrementally increasing weight every week by 5 gm increments up to 50 gm. The length of these sessions was increased in parallel until the animals did not exhibit any duress (i.e. pulling or biting the straps) while wearing the backpack. Significant Individual differences were evident in the duration of this habituation period, but all ultimately were habituated to wearing the backpack. While habituation to the backpack was necessary, the marmosets were minimally affected by the head-piece. In other words, marmosets rarely grabbed the hot mirrors used to track pupils. The only instance when this happened was a single occasion when the IR LED wire accidently slipped and was in a monkey’s peripheral view, and a separate occasion when an animal was chaired and a few drops of the liquid reward accidentally spilled on the hot mirror.

### 4. Multiple system synchronization

Our Neural recording systems, motion trackers and CEREBRO were all synchronized using an ESP32 microcontroller based custom board. CEREBRO and the motion tracking system sent TTL signals for different mode shifts that were recorded by the synchronization microcontroller. For synchronizing the neural signal, we sent a TTL pulse (100 mV |100 m) to the electrophysiology system (in recording mode) before putting it on the animal. The timestamp of this signal was logged by the sync microcontroller and since this signal was also logged into the real data stream of the electrophysiology system, we can easily align the time of this pulse in the data to other systems (CEREBRO and motion tracking). The board was connected to a computer via serial communication and the timing data was logged on to a console terminal (CoolTerm).

### 5. Electrophysiology

Neural activity was recorded with multi-electrode arrays (N-form, Modular Bionics, Berkeley CA) chronically implanted in V1. The N-form array has 64 channels in total embedded in 16 shanks located in a 4×4 grids evenly spaced 0.25 mm apart. Each shank has 4 iridium oxide electrodes located at 0.5, 0.375, 0.25, and 0 mm from the tip. Implantations followed the standard surgical procedures for chronically implanted arrays in primates. In each recording session, a wireless Neurologger (SpikeLog-64, Deuteron Technologies) was hosted in a 3.5 cm (width) x 2.5 cm (height) x 1.2 cm (depth) protecting case and connected to the array to record the extracellular voltage at 32 kHz. Spike sorting was performed offline using Kilo sort (60)and manually curated using the graphic user interface Phy.

### 6. Receptive field and tuning property of V1 neurons

For the receptive field mapping, marmosets were head-fixed and freely viewed the stimulus of 120 randomly positioned black and white flashing dots generated at a frequency of 10 Hz for 4 minutes. Eye calibration was performed offline with the GUI provided in this paper. We then used the calibrated eye position on the world camera images to estimate the visual input on the retina, and mapped the receptive fields using forward correction (61). For the orientation and spatial frequency tuning of the V1 neurons, we presented the full-field drifting gratings with 12 orientations (evenly spaced 30-degree intervals) and 3 spatial frequencies (0.5, 1, 2 cycles/degree). This information was used in calculating the tuning curves with the information from the screen; when calculating the tuning curves with the information from CEREBRO, we applied Hough transform on the world camera images to detect the lines and further calculated the orientation and spatial frequency of the gratings.

### 7. Head and body movement tracking using OptiTrack

The head and body movement of the animals were recorded using OptiTrack-an image-based motion tracking system (www.optitrack.com). For the head movements, the added 3 IR reflective beads (12mm in diameter) on the head-assembly of the animal. For the body, we added a single IR bead on to the backpack. The arena was fitted with 10 OptiTrack cameras strategically placed to cover every view of the arena. The cameras were calibrated before each session and sessions were performed only for calibrations where the error for the worst camera was less than 0.1 mm. The data collected from the session was manually curated to remove stray/false markers and any gaps were filled with linear interpolation.

### 8. Locomotion behavior analysis

Locomotion is detected with body and head tracking data from OptiTrack. We first calculated the velocity of the body IR bead and set a threshold of 5 cm/s (i.e. moving more than 20-30% the typical marmoset body length in under one second) after applying a 0.2 Hz low pass filter to select the period when the monkey had a fast body movement. However, not all the fast body movement came from locomotion; it could happen when the monkey made a body turn, or a sudden change of posture. We add two other conditions: 1) the duration of the period is larger than 3s; 2) the position of the head is lower than 18 cm. These two conditions are determined by manually matching the data with the video. After the locomotion period is detected, the rest of the time is defined as sedentary.

### 9. Freely moving eye calibration setup

Freely moving eye calibration paradigm consisted of a clear-plexiglass box (dimension: 30 cm x 30 cm x 40 cm). All sides of the box except the front panel were painted with black acrylic paint rendering it opaque. The front of the box was covered with IR long pass filter (OPTIR 1.0 NG 305 × 100, Newark) with a 10 cm x 10 cm window cut out in the middle at the height of 24 cm from the floor. The inside of the box had a wooden perch for subjects to sit on while looking out the window. The top of the box was lined with IR led strips (Near Infrared 850 nm NIR Single Chip DC12V 24W, 360DigitalSignage). The IR LEDs enabled video recording of subject’s behavior in the box since the IR dichroic allows the IR to pass through. Two synchronized FLIR cameras were mounted on the box. One camera was positioned to continuously monitor the subject’s behavior and the second camera was positioned to record the projection screen. A 2-axis galvanometric mirror driven LASER system (20Kpps speed Laser Galvo laser set, Wonsung Store) was installed on top of the box. This was used to draw various geometric shapes on the screen. All the shapes were generated in Python as plots with no axis. These plots were saved as SVG files and converted into ILD files using LASEROS software. The ILD files were loaded onto the galvanometric system using an SD card.

### 10. Error estimation for eye in world calibration

Eye/gaze data were calibrated (transformed to match the world scene) for a select number of trials. Following the initial process, the same transformations were applied to remaining trials. The eye/gaze position for these trials was overlaid on the world scene and exported as a video file. This video file was analyzed using a custom python script. Instances when the eye position was within 3.5 degrees of a visual feature and the residence time for the eye position for that location exceeded 60 ms were considered valid instances of looking at the feature. For such valid instances, the closest distance (radius of the smallest circle that touches the visual feature with eye position as the center) between the eye position and the closest visual feature (in degrees) was used as the estimate of the error associated with our eye calibration. This places an upper bound on the error as the animal’s actual eye position would not exactly match the center of the object even when viewing it due to the object’s extent, as well as fixational eye movements and drift.

### 11. Camera and communication interface

OV4689 camera modules with two MIPI lanes and customized Flex cable length of 15 cm (SincereFirst, Guangzhou, China) was used both as the eye camera and the world camera. The cameras operated on a serial interface called MIPI CSI-2 (Mobile Industry Processor Interface Camera Serial Interface). Since microprocessors are the core of the system a STMIPID02 MIPI CSI-2 deserializer was used to deserialize CSI communication from a MIPI CSI-2 camera sensor to DCMI and improve the temporal resolution of the incoming data.

### 12. Printed Circuit Board and data storage

Our custom PCBs use STM32H750 which is a 32-bit Arm Cortex-M7 core processor running up to 480 MHz. It is better suited for interfacing with camera sensors since it embeds a digital camera interface (DCMI). Each PCB can be programmed and debugged using the Serial Wire Debug (SWD) mode. The PCBs store the raw camera data on a locally mounted, UHS-1 (Ultra High Speed) class, SD card. The binary files are decoded offline to get an AVI format video of the stream and a CSV file with timestamp and IMU data. Two PCBs are stacked on top of each other and are enclosed in a custom designed 3D printed casing with a custom designed back harness. To measure camera synchronized movements (acceleration and orientation) of head, the boards interface with an inertial measurement unit (IMU). The PCB boards also feature a 0.96“ SPI TFT display which allows the users to preview the captured camera image and ensure the placement of the head-piece at the start of the session.

