## Supplementary information for "Active vision in freely moving marmosets using head-mounted eye tracking"

### Supplementary Information 1: Detailed description of the head assembly

The head assembly is held on the animal’s head using two 3D printed horns that are permanently glued to the animal’s head post with the help of dental cement/ orthodontic acrylic resin. (*fig 1c*). All the parts are printed with Titanium (Ti-6-Al-4V) using Direct Metal Laser Sintering (DLS). All the 3D parts are held together with a 2 mm (diameter) stainless steel rod curved to match the profile of the animal’s forehead (*Supplementary fig 1a* shown in yellow). General assembly guidelines are shown in *Supplementary fig 1b.*

| **a)**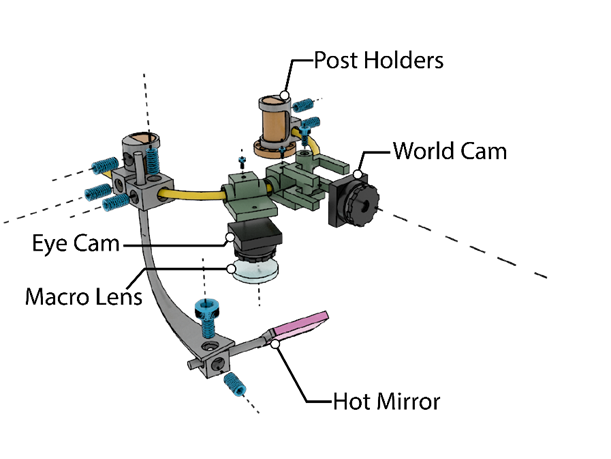 | **b)**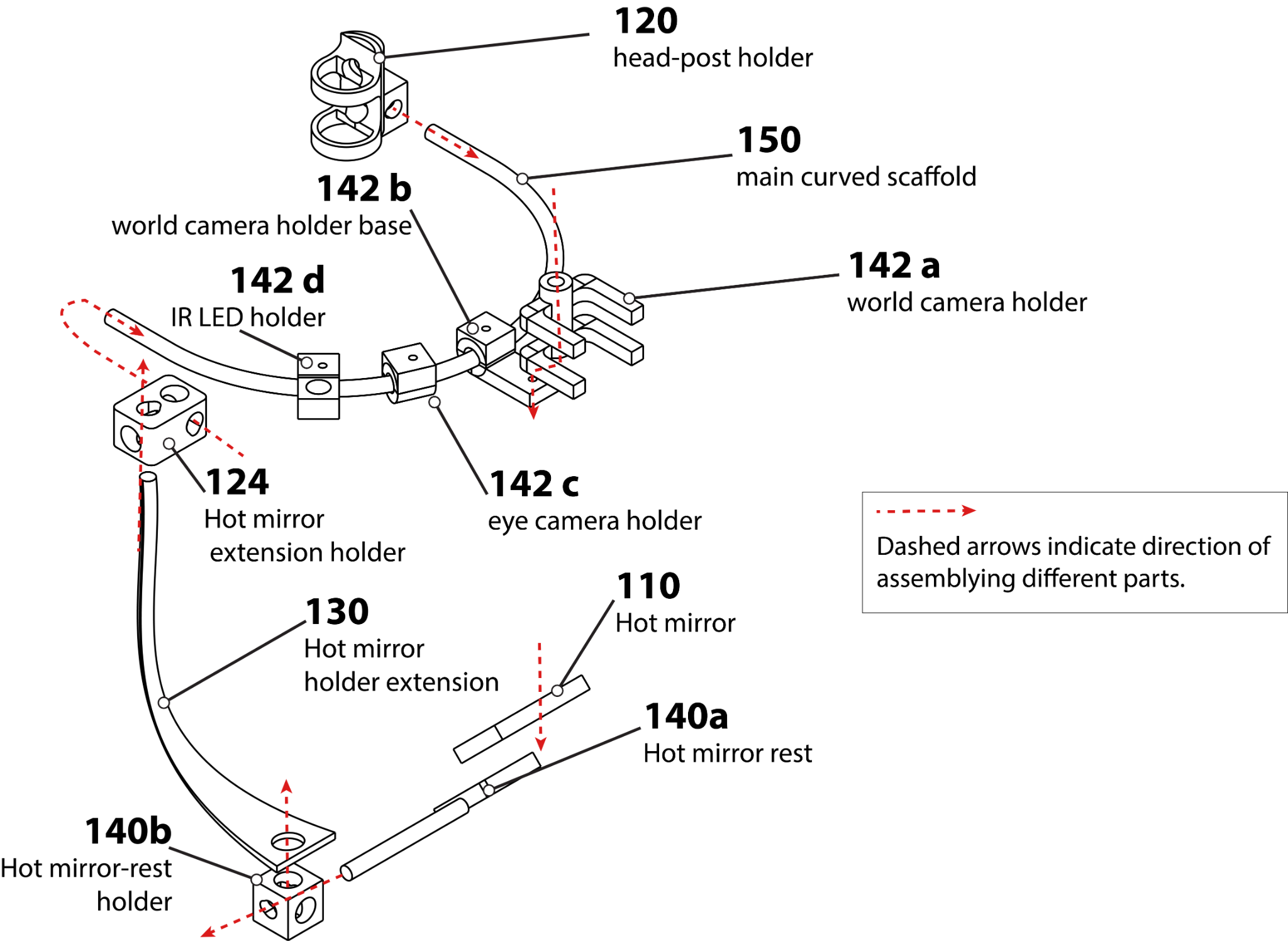 |
| --- | --- |

**Supplementary Figure1:** **a)** 3D render of the head assembly. **b)** Assembly guidelines for putting together all the pieces of the headpiece.

The key anchoring structures in the head assembly are the head-post holders (marked *120* in s*upplementary fig b*). The headposts are structurally the female counterparts to the headposts on the animal's head. They latch to the head-posts with the help of a M4 set screw. The main scaffold i.e. a curved steel rod (2mm in diameter, marked as *150*  in *supplementary fig 1b*) passes through the headpost holders and all the other parts are directly or indirectly attached to this curved scaffold. The eye camera is attached to the eye-camera-holder (*142c in supplementary fig 1b*) using fast curing orthodontic acrylic resin.The camera holder can rotate along the axis of the curved scaffold and is fixed at the final position with a M2 screw. Right next to the eye-camera-holder is the IR LED holder (*142d*) that holds a 3mm 940 nm IR LED. This LED is connected in parallel to the battery on the backpack. The world camera is attached to the scaffold indirectly via 2 connected pieces. There is a small piece (world camera holder base; *142b*) that attaches to the scaffold and the other piece (world camera holder; *142a*) holds the world camera. Both the pieces use M2 screws for gripping the structural piece. One of the crucial pieces of the head assembly is the hot mirror which has to be positioned strategically and requires a design that allows for free adjustment of the mirror at the desired position. This is achieved by using a set of 4 small pieces that indirectly attach the hot mirror to the main scaffold. There is a long extension (*130*) that can slide in and out of a small piece (*124*) which in turn holds the hot mirror submodule to the scaffold with M4 set screws. The sliding of the extension rod allows for adjustment of the height of the hot mirror. A 45-degree 12.5 mm x 12.5mm hot mirror (Edmund optics) is glued to the hot mirror rest (*110*) which is attached to the extension rod via a small cube piece (*140b*). The cube allows the rotation of the hot mirror and the hot mirror rest can also be adjusted for how far the mirror needs to be placed for best view of the eye.

### Supplementary Information 2: Synchronization of various systems using ESP32 microcontroller.

In order to synchronize the various systems that are running on different clocks, we used the approach of synchronizing every system to a common clock. We used an ESP32 microcontroller as the synchronizing clock.We refer to the finished design with all the passive electronics as SyncBox. CEREBRO is capable of sending a TTL pulse for every button press of the user. This signal is sent to the sync box and it triggers a hardware interrupt. The timing of this interrupt is logged to a serial terminal on the computer. Our motion tracking system (Optitrack) can also send a TTL at the start and stop of a recording session. This TTL was also logged in the same fashion on the serial monitor. For synchronization of the neural recording device, we used a different approach. The SyncBox can also send a TTL out on a button press. We connect the output of a 100ms, 100mV TTL to our electrophysiology recording device. This shows as a pulse signal in our raw data and the timing for it is logged by the SyncBox.

**a)**
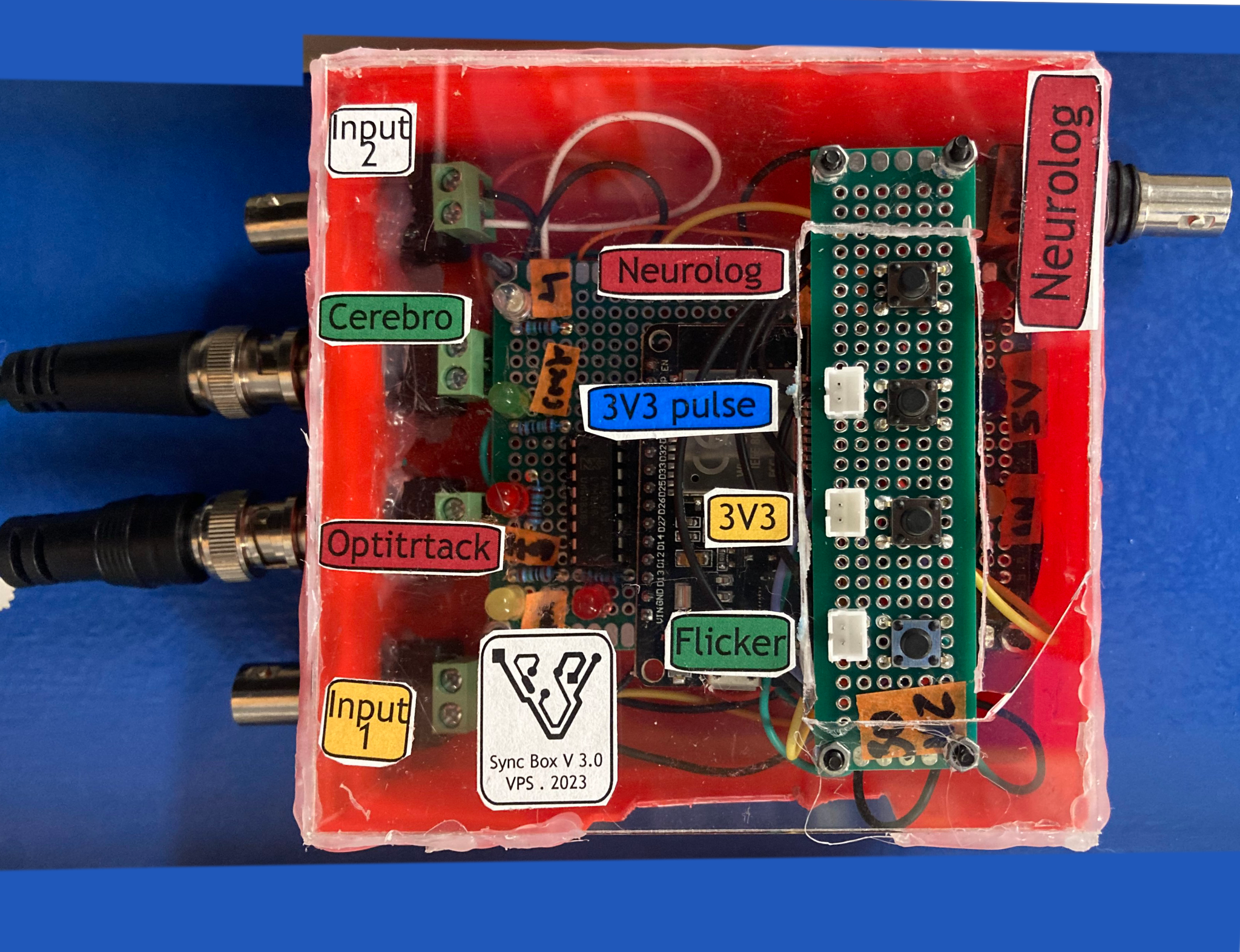

**b)**

**b)**
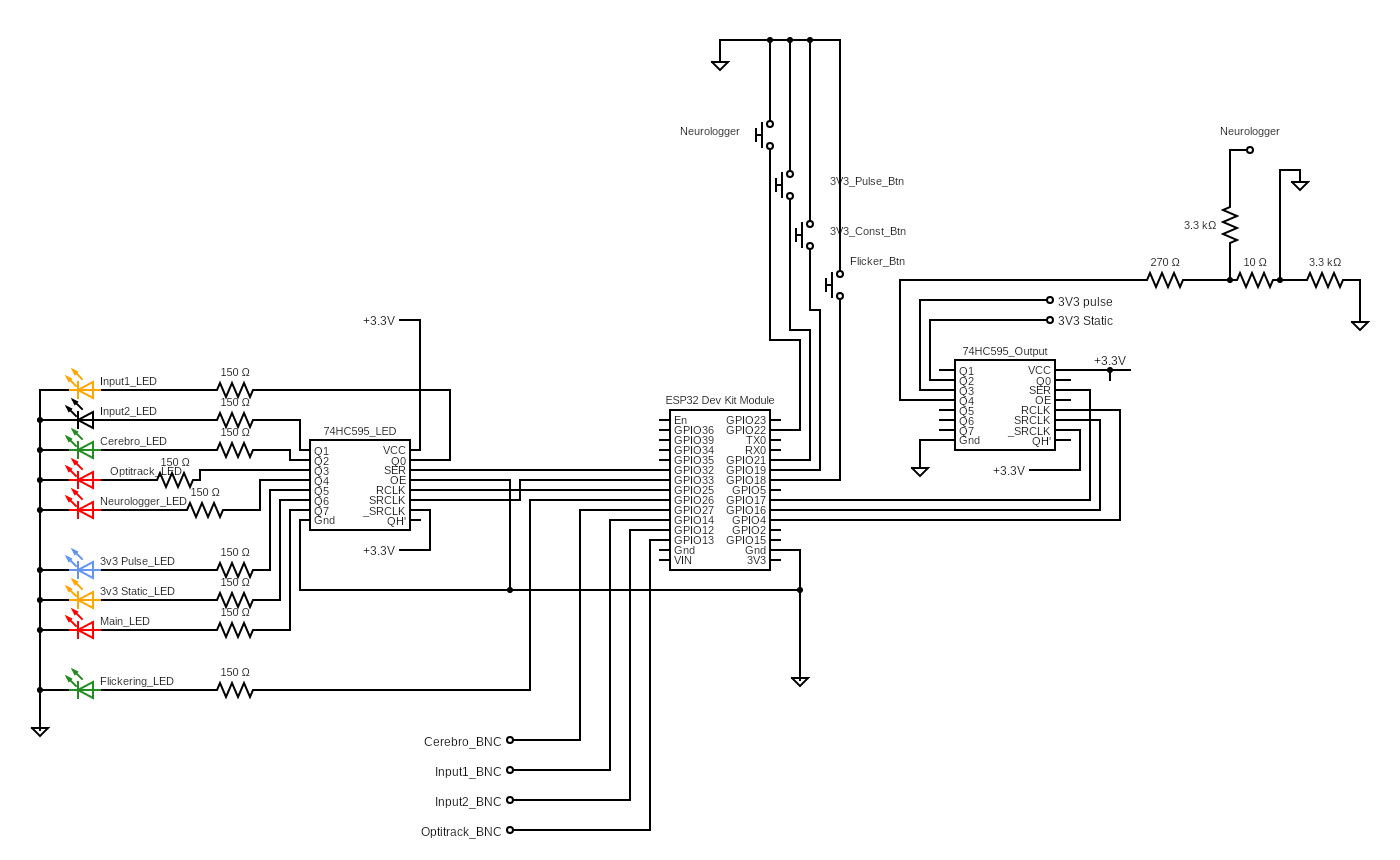

**Supplementary Figure2: a)** Sync Box for synchronizing different devices. **b)** Schematic of the SyncBox.

### Supplementary Information 3: Flow of information in the Custom PCB.

The PCB has several peripherals. It has connectors for the camera, IMU, SD card, GPIO pins and battery, LCD screen and push buttons to allow for resetting the board and to receive user input to help it switch modes (*Supplementary fig 3a*). Supplementary figure 3b shows the flow of data through the board. The microcontroller unit (MCU) receives data from the camera and IMU (1 and 2). The data from the camera is first converted to the right format using the MIPI deserialized chip. MCU then stores the data to the SD card during recording phase (5) and to the LCD screen during preview mode (6).

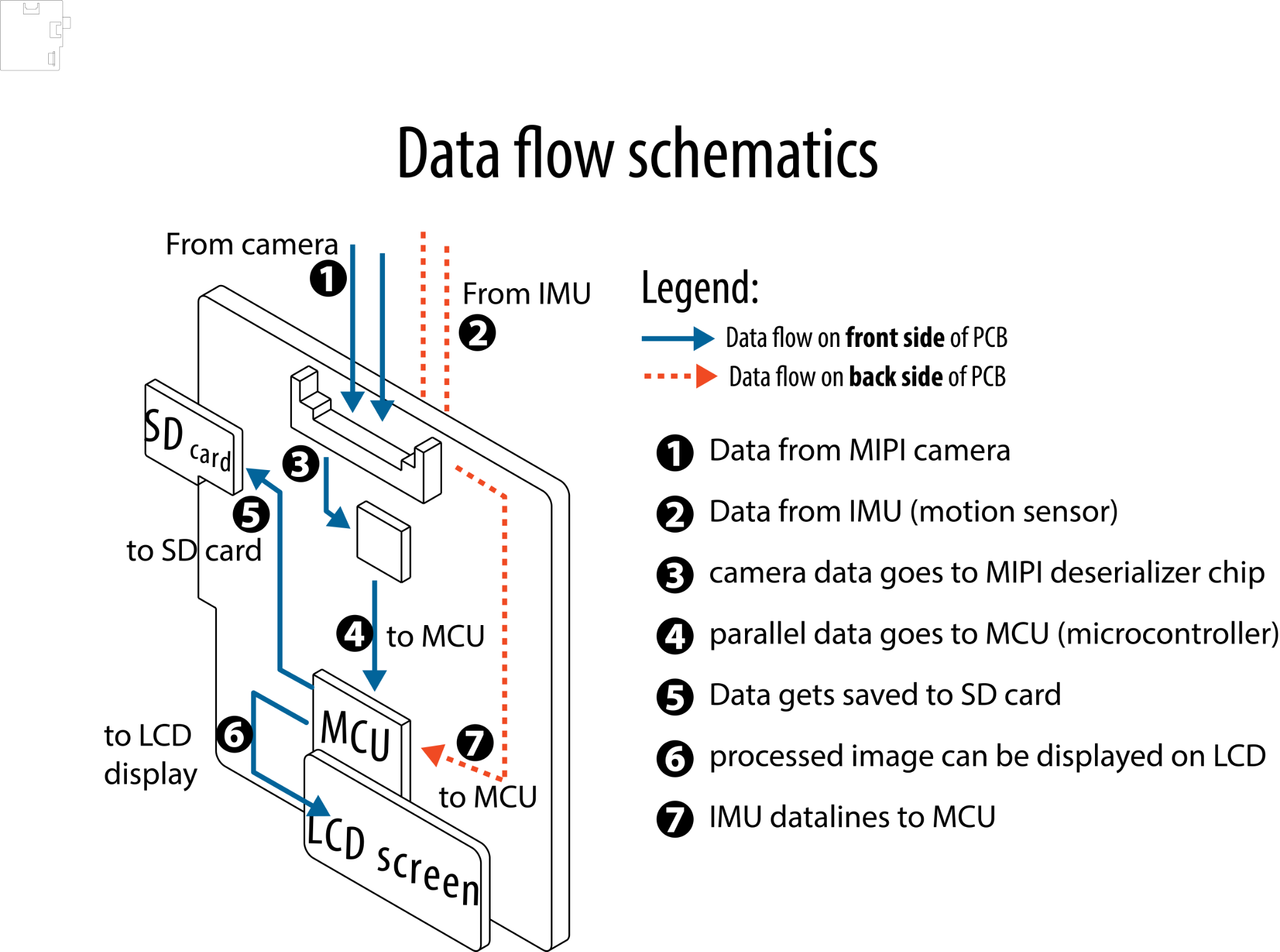

| **a)**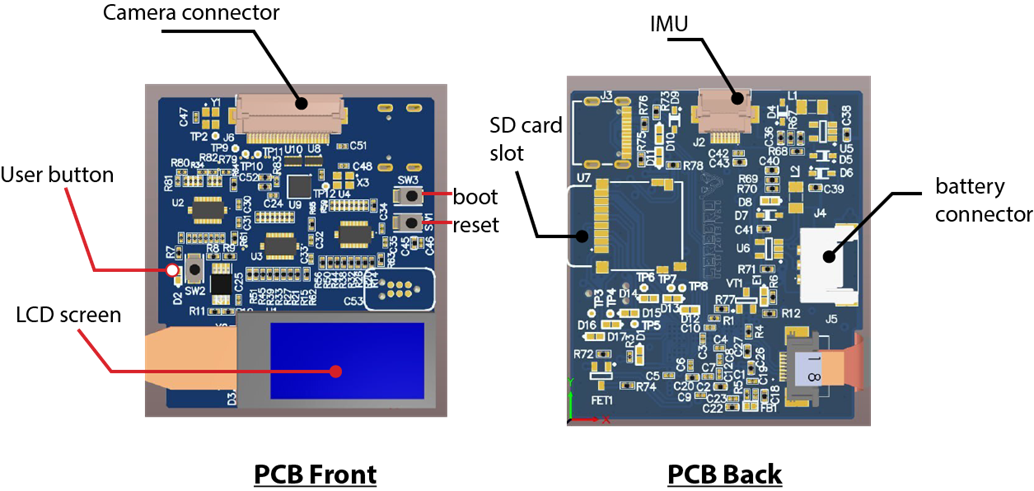 | **b)** |
| --- | --- |

**Supplementary fig3: a)** front and back of the custom designed PCB. **(b)** flow of information through the PCB board.

### Supplementary Information 4: Segmentation Artificial Neural network for pupil detection

Since the thresholding approach has several caveats in a freely moving animal, we switched to an ANN approach. We trained UNET,a segmentation Neural network, to crop the region of interest that can be followed by thresholding and ellipse fitting for pupil detection. Whereas classical convolutional networks are classifiers which output a single class label with a confidence value, the UNet architecture achieves this by assigning a class label to each pixel of the image. The core logic for the UNet implemented here was to supplement a classical convolutional network (contracting path) with successive upscaled layers such that the final output image is of the same dimension with object of interest segmented out as white (>0) and background set to black (0). The UNET implementation consists of 3 main steps: a) preparing the dataset and training the UNET; (b) implementing the UNET to segment IRIS (c) pupil detection after thresholding and ellipse fitting.

#### a) Preparing the training dataset: We prepare the dataset by having 2 main folders. One of the folders consists of images from the CEREBRO system and the other folder contains corresponding masked images with the iris of the eye being white (object of interest) and the rest of the image black (background elements) (*supplementary fig 4a*). We use this dataset to train the model using Tensorflow libraries in python.

#### b) UNET implementation: We implemented the segmentation of eye videos by bundling the neural network into a python based graphic user interface (*supplementary fig 4b*). The executable file for the GUI is available at: <https://github.com/Vickey17/UNET_GUI_exe>. The users have an option of loading the “*raw eye timestamps*” and adjusting the orientation of the video frames. Upon execution of the GUI, a new video is created with the same name and “_UNET_Result” appended to it. The progress of the process can be tracked with the progress bar at the bottom.

#### c) Pupil detection: The segmented video from the UNET is read in MATLAB and we perform blob detection (https://www.mathworks.com/help/vision/ref/vision.blobanalysis-system-object.html) to detect the pupil from every frame. During blob detection, there is a possibility that there are some stray blobs in the image. We only log the largest blob detected in the image as pupil while all the stray blobs are discarded. The center of the detected blob is taken as the pupil.

| **a)**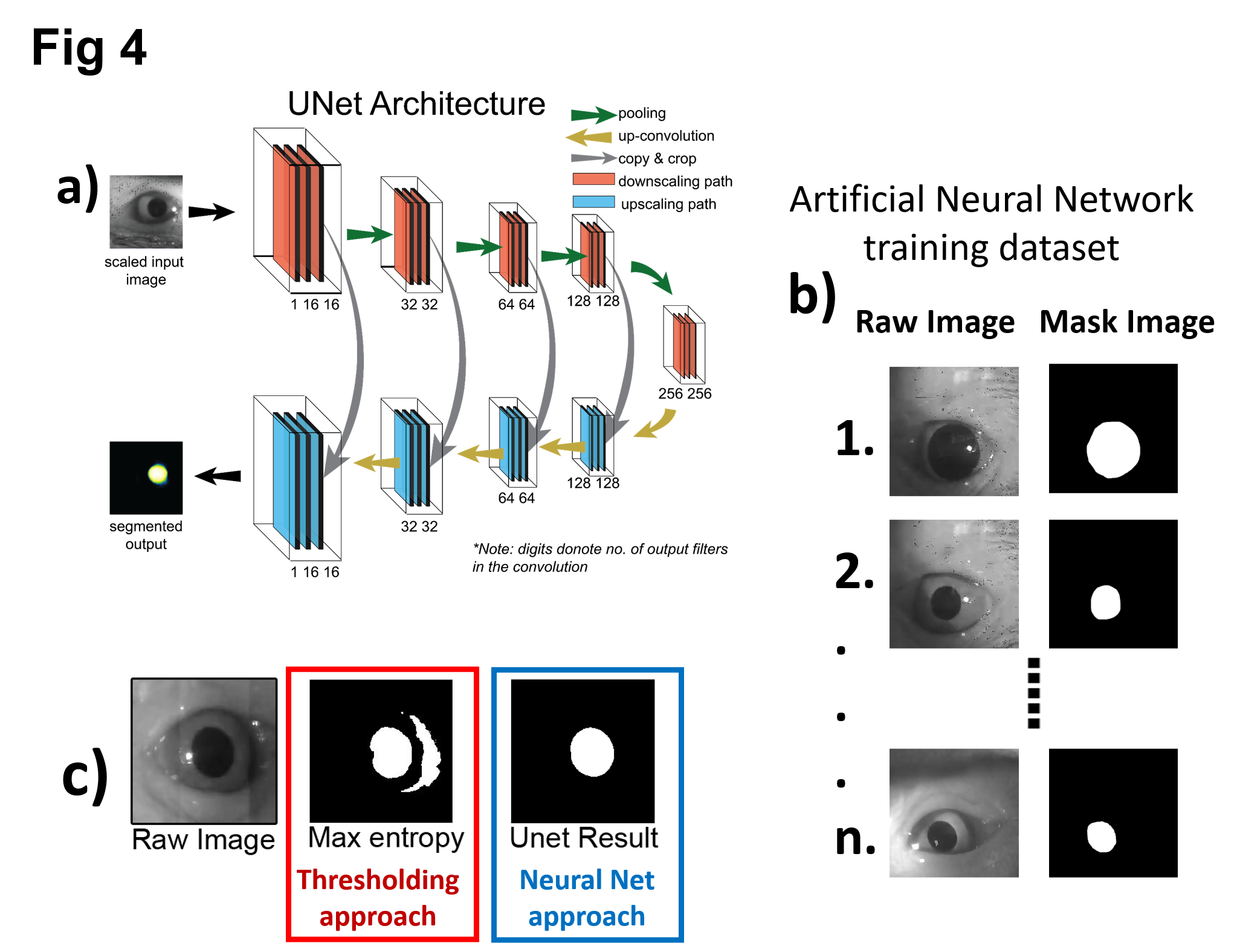 | **b)**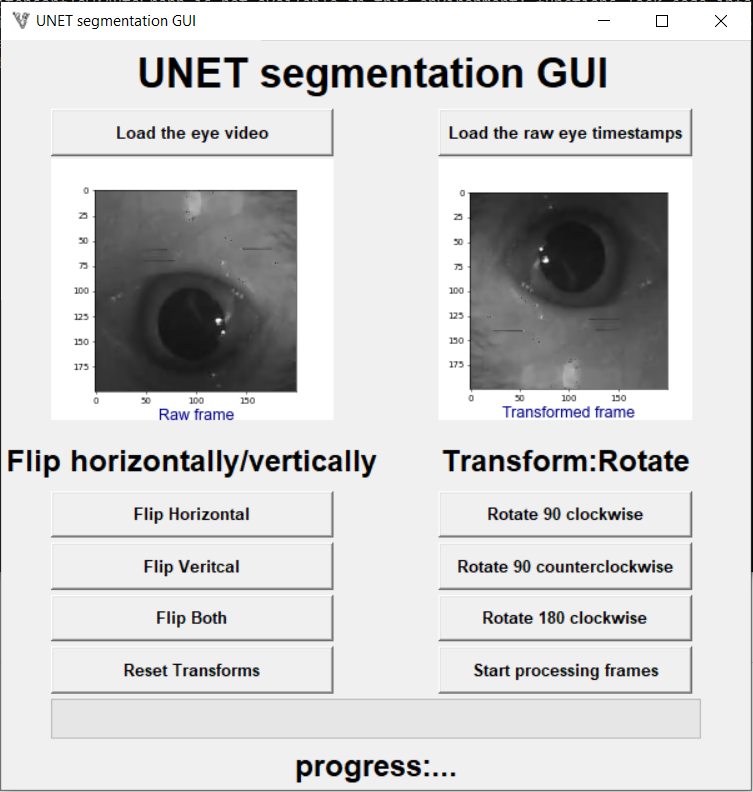 |
| --- | --- |

#### **Supplementary figure 4 a)** Illustration for the training dataset preparation. Each image has a masked counterpart with the same name stored in a separate folder. The mask allows the ANN to learn features of interest from the real image. **b)** UNet implementation GUI. The GUI allows users to open an image captured using CEREBRO and save a segmented video.

### Supplementary Information 5: Comparison between RITNET and UNet described in this study.

We compared the performance of UNet described in our study with RITNET described by (Chaudhary et al. 2019) on human eye data (*Suppl. fig. 5a*) for which RITNET was pre-trained while our network was not. Pupil detection from UNet is marked with a red bounding box while that from RITNET is shown in yellow. We observed that RITNET was clearly superior to our UNet in this case but for marmoset eyes that have significantly different visual features, our UNet performed better for both trained (*Suppl. fig. 5b*) and untrained marmoset eye images (*Suppl. fig. 5c*).

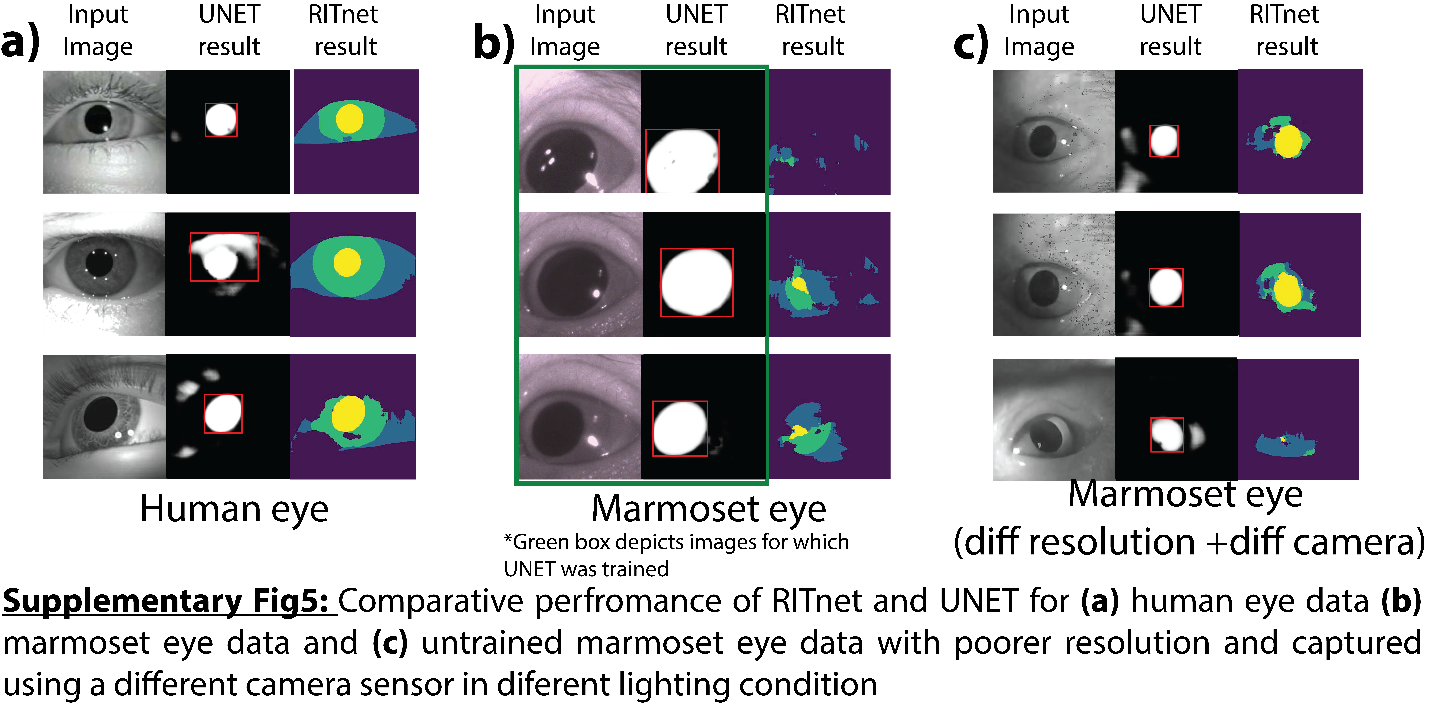

#### **Supplementary figure 5:** Comparative performance of RITnet and UNET for **(a)** human eye data **(b)** marmoset eye data and **(c)** untrained marmoset eye data with poorer resolution and captured using a different camera sensor in different lighting condition

Supplementary Information 6: Gaze statistics plotted for each animal separately.

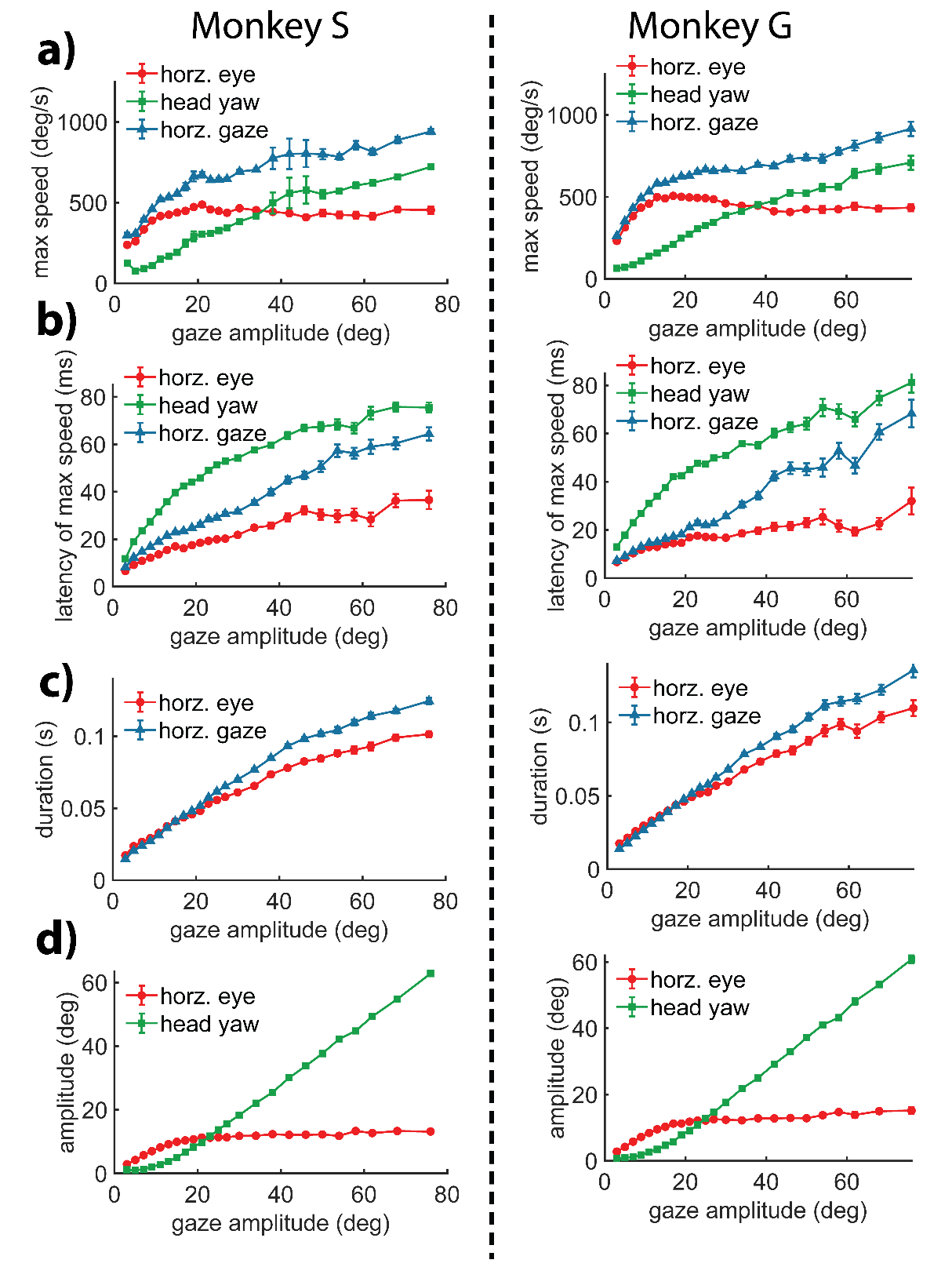

**Supplementary figure 6:** Gaze statistics plotted for each animal separately **(a)** max speed **(b)** latency **(c)** duration and **(d)** amplitude as a function of gaze amplitude.

Supplementary Information 7: Eye behavior statistics plotted for each animal separately.

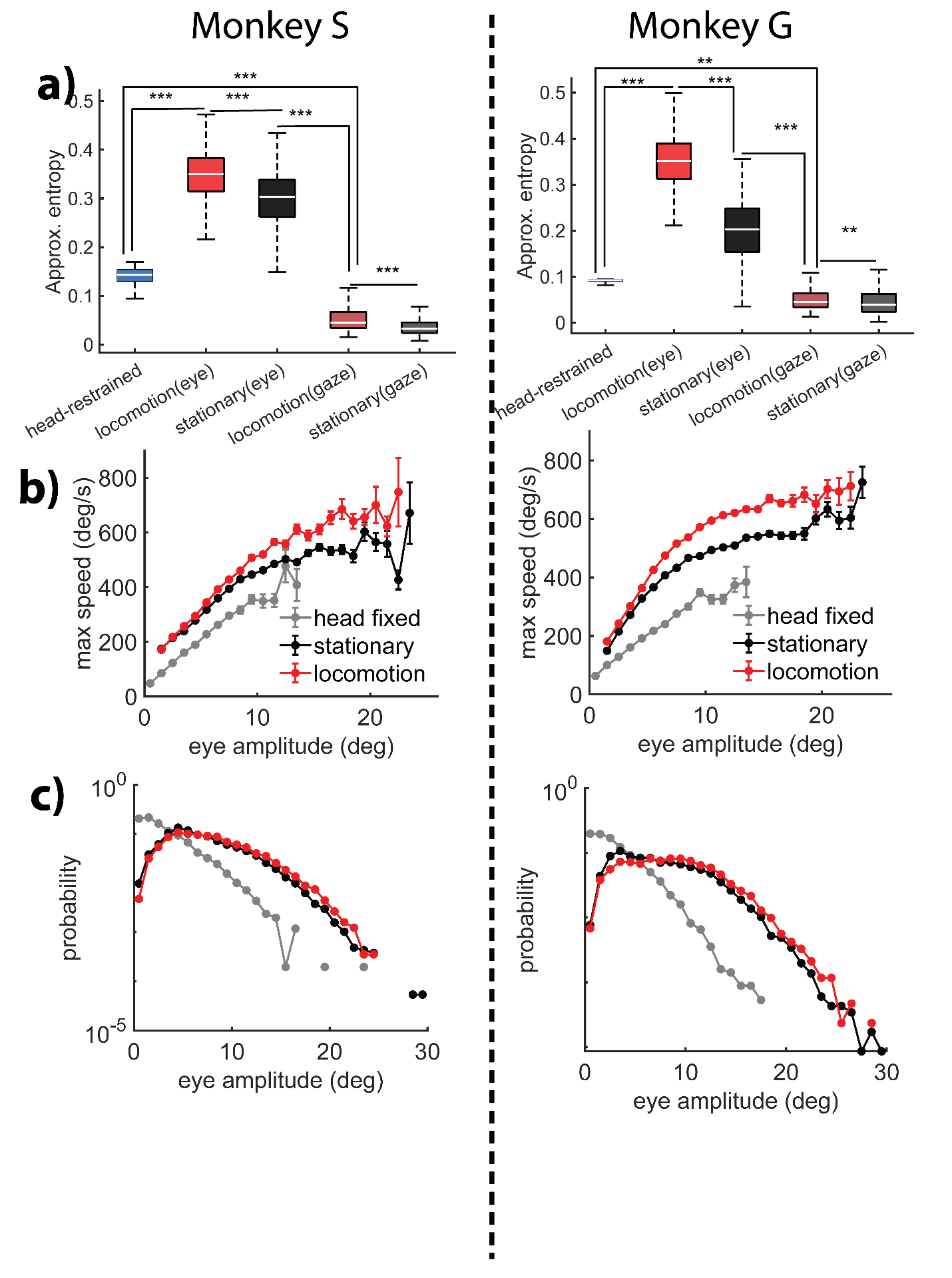

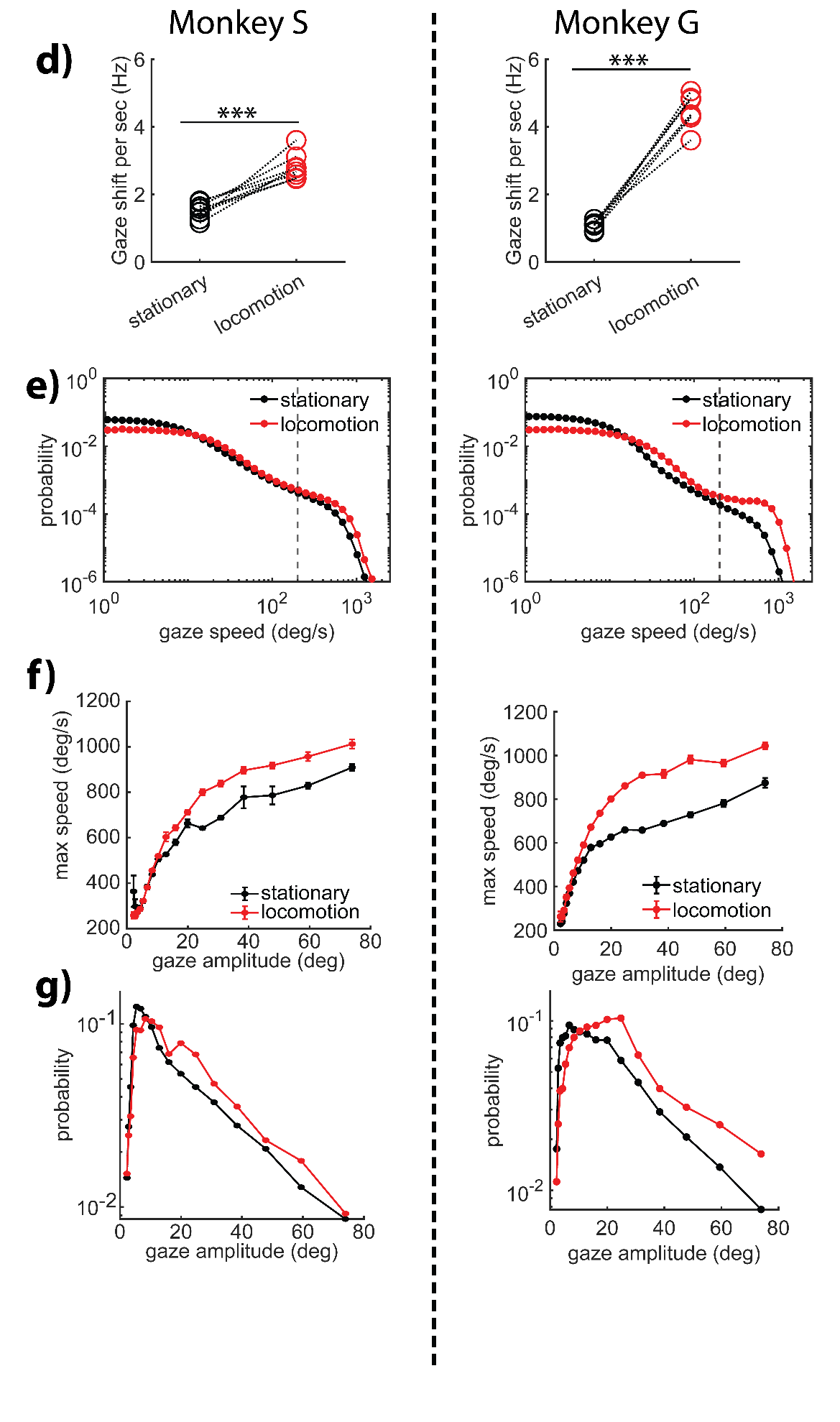

**Supplementary figure 7:** Eye behavior statistics plotted for each animal separately **(a)** approximate entropy for head restrained and freely moving conditions for eye and gaze. **(b)** max speed as a function of eye amplitude. **(c)** distribution of eye amplitude in stationary vs locomotion context. **(d)** frequency of saccades in locomotion vs stationary. **(e)** distribution of gaze speed across different behavioral context. **(f)** max speed as a function of gaze amplitude for stationary vs locomotion. **(g)** distribution of gaze amplitude in locomotion vs stationary epochs.

Supplementary Information 8: Eye contribution to RMS stabilization.

The difference between head and gaze RMS stabilization gave us the contribution of eye movements to the gaze RMS stabilization. We observed that the eyes’ contribution to stabilization is increased during locomotion as compared to stationary behavioral epochs.

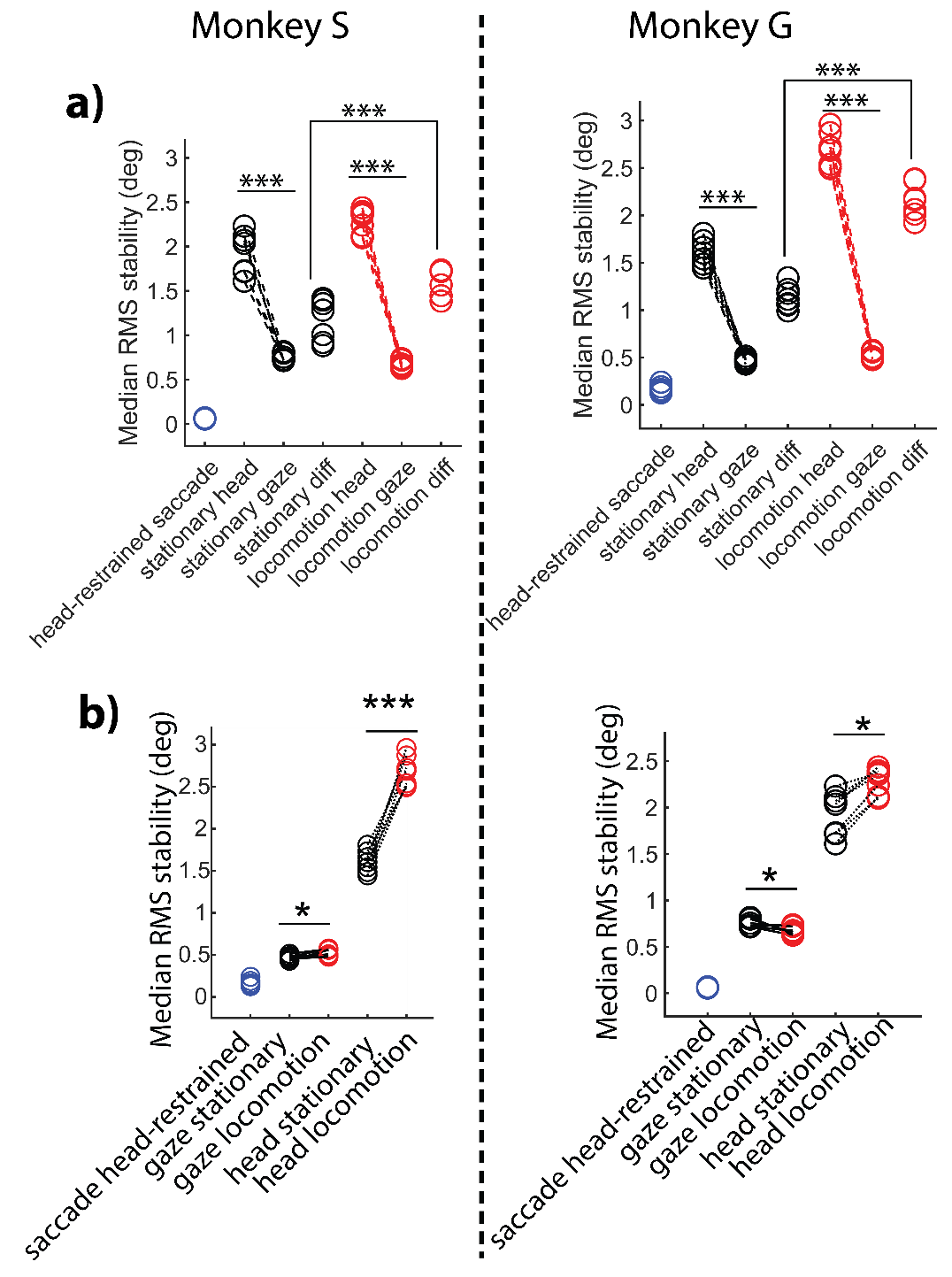

**Supplementary figure 8:** Eye contribution to RMS stabilization. **(a)** eyes contribute significantly more to RMS stabilization during locomotion vs stationary behavior (locomotion diff vs stationary diff in the plot). **(b)** Head stability reduces drastically during locomotion, but gaze stability improves for Monkey S and slightly decreases for Monkey G.

Supplementary Information 8: Gain in VOR during locomotion

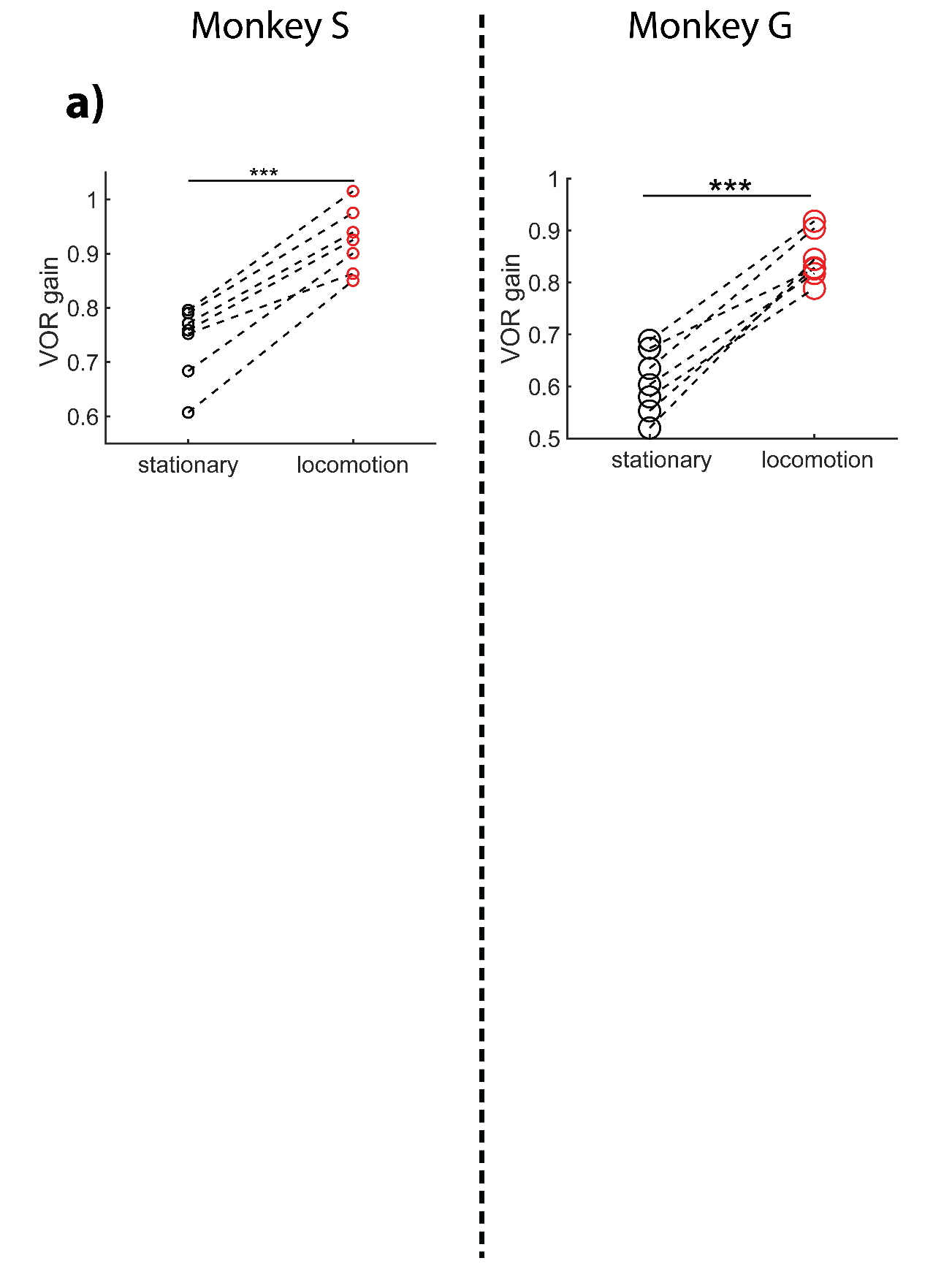

**Supplementary figure 8:** Gain in VOR during locomotion

Supplementary Information 9: Duration of gaze shifts in head-restrained and freely moving conditions

Gaze shifts / saccades are slower in head-fixed context as compared to freely-moving conditions.

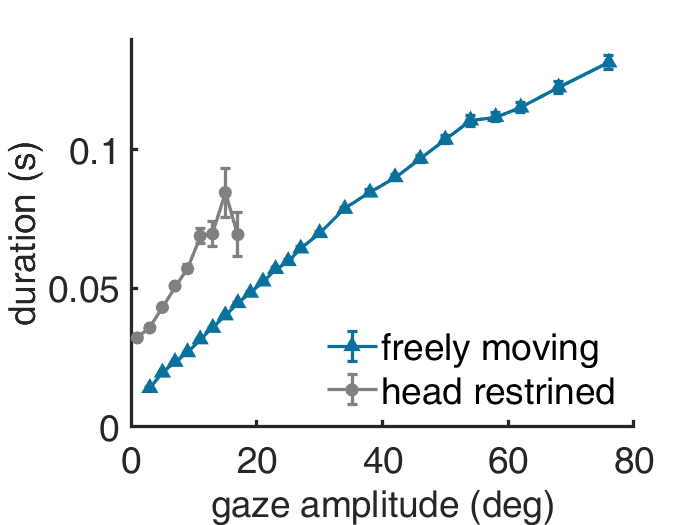

**Supplementary figure 9:** Duration of gaze shifts in head-restrained and freely moving conditions
